# De novo design of CR2 binder as vaccine scaffold

**DOI:** 10.64898/2026.08.28.747755

**Authors:** Yuan-Tao Liu, Yi-Na Yin, Hui-Qin Xu, Wen-Ting Du, Chu Xie, Pei-Huang Wu, Hang Zhou, Bing-Zhen Cheng, Guo-Long Bu, Guo-Kai Feng, Qian Zhong, Zheng Liu, Mu-Sheng Zeng, Cong Sun

**Author notes:** These authors contributed equally to this work (Yuan-Tao Liu, Yi-Na Yin, Hui-Qin Xu, Wen-Ting Du). Corresponding authors (C.S); (M.S.Z.); (Z.L.).

## Abstract

Efficient B cell activation during vaccine-induced humoral immunity relies on both B cell receptor (BCR) antigen recognition and synergistic signaling from co-receptors. Complement receptor 2 (CR2), the primary BCR co-receptor on B cells, lowers the activation threshold and amplifies downstream kinase signaling by orders of magnitude when engaged by complement fragment C3d-decorated antigens. Targeting CR2 therefore represents a rational vaccine enhancement strategy, yet native C3d suffers from low affinity, poor stability, and manufacturing challenges. Here, we report the de novo design of a highly stable, high-affinity CR2 binder using deep learning–driven protein design methods. Biophysical characterization, high-resolution cryo-EM structural determination, and functional assays *in vitro* and *in vivo* confirm that the designed binder matches computational design models and specifically engages CR2 to boost B cell activation. When fused to antigen as a vaccine scaffold, the trimeric CR2 binder elicits robust humoral immune responses comparable to nanoparticle vaccines, while retaining the simplicity of single-chain protein production. Our work establishes a modular CR2-targeting vaccine scaffold platform with broad translational potential for next-generation protein vaccines.

## Introduction

Induction of antigen-specific antibody production via B cell activation is the central mechanism of most prophylactic vaccines (*1–3*). Enhancing B cell antigen recognition and activation is therefore a core strategy to improve vaccine efficacy, yet current approaches rely heavily on empirical adjuvants, with few options for antigen-specific B cell targeting (*2*, *4*). Adjuvants improve immunogenicity nonspecifically, but they do not direct antigen to B cells or engage the co-receptor complexes that set the B cell activation threshold(*5*, *6*). A vaccine platform that combines antigen-specific B cell targeting with co-receptor engagement would address a gap that adjuvant-only strategies cannot fill(*3*, *7*). Several antigen-linked B-cell-targeting strategies have been explored. Targeting antigen to CD19 directly recruits the signaling component of the B-cell co-receptor complex but requires antibody-derived delivery modules(*8*), whereas targeting CD180(*9*) or fusing antigen to BAFF, APRIL or CD40L(*10*) engages distinct and often pleiotropic activation or survival pathways. Conversely, engagement of inhibitory co-receptors such as CD22(*11*) or FcγRIIB has been exploited to induce antigen-specific tolerance rather than protective immunity(*12*). CR2 is distinct in that its natural ligand C3d couples antigen recognition directly to the CR2-CD19-CD81 complex, providing a physiologically defined mechanism to lower the BCR activation threshold and motivating our focus on CR2(*12*, *13*).

Optimal B cell activation requires coordinated signaling from the BCR and cell-surface co-receptor complexes(*14*). The CR2-CD19-CD81 complex is the dominant activating co-receptor on B cells. Its engagement reduces the BCR activation threshold by 100- to 10,000-fold and amplifies downstream kinase signaling cascades(*15–18*). During natural humoral responses, complement component C3b covalently attaches to antigens and is proteolytically processed to C3d, which engages CR2 on B cells(*19*, *20*). Co-ligation of BCR (by antigen) and CR2 (by antigen-bound C3d) drives robust activation, survival, and clonal expansion of antigen-specific B cells(*16–18*, *21*). This natural co-engagement mechanism makes CR2 a long-recognized but unrealized vaccine target.

Native C3d has been explored as a molecular adjuvant to boost vaccine immunogenicity(*13*, *19*), but its clinical translation is hindered by critical limitations. Monomeric C3d binds CR2 with only micromolar affinity, insufficient to break tolerance in anergic B cells(*13*, *22*). Additionally, native complement fragments are difficult to express and purify at scale, exhibit limited conformational stability, and carry risks of off-target complement pathway modulation and autoimmune sequelae(*23*–*25*). Developing a stable, high-affinity CR2-targeting platform has remained an open problem for over three decades, primarily because native C3d’s micromolar affinity demands tandem multimerization for biological effect, and any sequence homology to host complement risks breaking B cell tolerance to self. A non-native CR2 binder with high affinity, manufacturable stability, and no complement cross-reactivity would resolve both limitations simultaneously.

Recent advances in deep learning–based computational protein design have enabled de novo generation of proteins with predefined architectures and binding interfaces(*26–28*), providing new opportunities for engineering functional protein scaffolds(*29*, *30*). Here, we combine de novo binder design with symmetric oligomerization engineering to generate a CR2-targeting protein scaffold. The trimeric CR2 binder engages B cell CR2 with nanomolar apparent affinity, mimics the clustered geometry of complement-opsonized antigens, and can be fused to antigen as a single polypeptide chain. We use biophysical, structural, and *in vivo* assays to test whether this scaffold enhances B cell activation and humoral immunity relative to native C3d fusion and self-assembling nanoparticle display. The work establishes a generalizable framework for CR2-targeted vaccine engineering and provides a route to convert a longstanding immunological mechanism into a manufacturable protein module.

## Results

### De novo design and optimization of CR2 binders

CR2 specifically recognizes C3d-opsonized antigens via its N-terminal short consensus repeat domains 1 and 2 (SCR1-2), a molecular interaction that underlies the co-receptor’s ability to lower B cell receptor (BCR) activation thresholds(*31*). Direct engineering of C3d to improve affinity is constrained by its structural complexity and the risk of disrupting endogenous complement homeostasis(*32*, *33*). We therefore sought to design a de novo protein that recapitulates CR2 binding activity while offering improved stability, affinity, and safety (Fig. 1A).

**Fig. 1.**
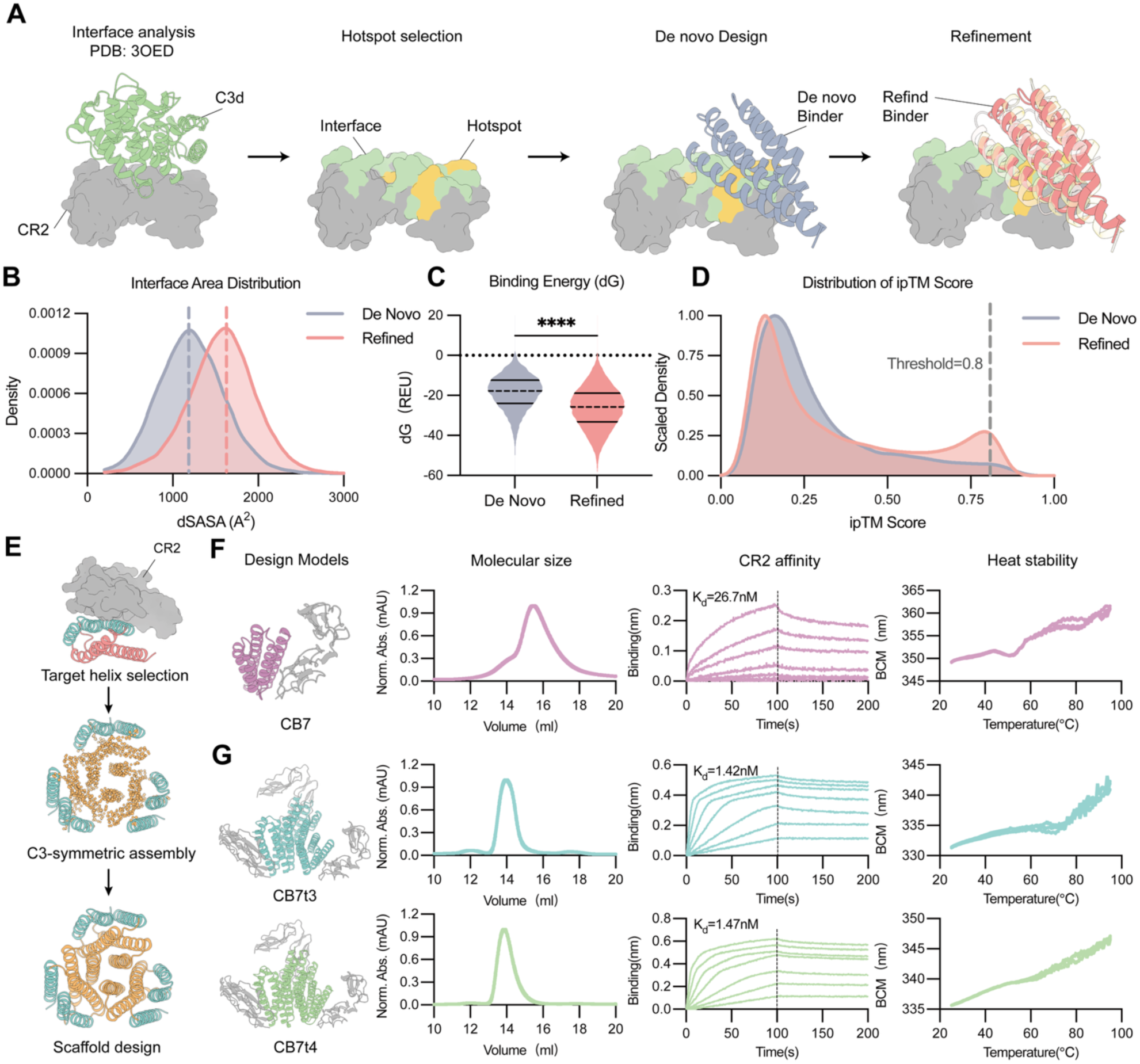
De novo design and characterization of monomeric and trimeric CR2 binders. (A) Schematic of the de novo CR2 binder design workflow, including CR2–C3d interface analysis, hotspot residue definition, de novo backbone generation, and partial-diffusion affinity optimization. (B) Kernel density plots of buried solvent-accessible surface area (ΔSASA) for initial de novo designs and refined binders. Optimization shifts the distribution toward higher ΔSASA, indicating tighter interface packing. (C) Binding free energy (ΔG, Rosetta energy units) distributions before and after partial-diffusion refinement. Refined designs are enriched in lower-energy states, consistent with improved thermodynamic stability. Dashed line, Q2 (median); solid lines, Q1 and Q3. Statistical significance was assessed using a two-sided Welch’s t-test. \*\*\*\**P* < 0.0001. (D) Scaled kernel density plots of Chai-1 interface predicted template modeling (ipTM) scores for initial and refined binder designs. The y-axis represents scaled density. Refined binders are enriched in high-confidence models (ipTM > 0.8), indicating improved interface reliability. (E) Design strategy for C3-symmetric trimeric CR2 binders. The CR2-binding helical motif (cyan) is arranged symmetrically around the C3 axis and scaffolded into a trimeric assembly with a de novo– designed hydrophobic core (orange). (F) Biophysical and kinetic characterization of the monomeric CR2 binder CB7, including size-exclusion chromatography (SEC) profile, binding kinetics to human CR2 measured by biolayer interferometry (BLI), and thermal stability measured by differential scanning fluorimetry (DSF). (G) Biophysical and kinetic characterization (apparent *K*_d_, avidity-inclusive) of trimeric CR2 binders CB7t3 and CB7t4, including SEC profiles, binding kinetics to human CR2 measured by BLI, and DSF thermal stability. Data in (F), (G) are sequential signal data from an individual measurement. One sample data is presented for molecular size and CR2 affinity measurement and triplicate sample data are presented for heat stability measurement.

We first delineated the C3d–CR2 SCR1-2 interaction interface using Rosetta InterfaceAnalyzer(*34*) and PDBePISA(*35*), identifying key polar/electrostatic residues (S38, D112, R48, K61) and hydrophobic interface residues (Y36, V46) that mediate binding. Specifically, Y36 forms a critical side-chain hydrogen bond to stabilize interface packing, while V46 reinforces the complex via backbone hydrogen bonds and hydrophobic stacking. Leveraging these insights, we defined Y36 and V46 as primary hotspot constraints and incorporated adjacent surface residues (L57, L64, I66) as spatial constraints to orient the binder at the SCR1–SCR2 domain junction (Fig. S1A).

We used RFdiffusion(*26*) to generate protein backbones conditioned on these constraints, followed by sequence design with ProteinMPNN(*27*) and structural validation with Chai-1(*36*). This first-round pipeline yielded 12,000 candidate CR2 binders (Fig. S1B). To further improve binding affinity, we performed partial diffusion optimization on the top 5 designs (ranked by binding free energy), generating 4,000 candidates per design for a total of 20,000 optimized variants(Fig. S1C).

Computational analysis confirmed the efficacy of our optimization strategy. The distribution of buried solvent-accessible surface area (ΔSASA) shifted significantly rightward after optimization, indicating tighter interface packing and larger contact areas (Fig. 1B). Concurrently, calculated binding free energy (ΔG, Rosetta energy units) shifted to lower energy states, consistent with improved thermodynamic stability (Fig. 1C). Importantly, structure prediction with Chai-1 revealed a marked enrichment of optimized designs within the high-confidence regime (ipTM > 0.8), indicating that the designed sequences fold into well-defined, native-like binding interfaces (Fig. 1D). Based on a composite ranking of binding energy and structural confidence, we selected 12 top-performing candidates for synthesis and experimental characterization. Biolayer interferometry (BLI) revealed that 3 designs exhibited specific CR2 binding activity with cross-reactivity to both human and murine CR2. The strongest binder, designated CB7, was selected for further engineering and characterization (Fig. S1D).

### Trimerization of the lead CR2 binder enhances affinity and stability

Natural C3d fusion antigens have limited immunogenicity as monomers, and multimeric tandem fusions are required for robust B cell activation(*13*, *37*). Additionally, many viral surface antigens adopt trimeric conformations, and multivalent antigen display is a well-established strategy to enhance B cell recognition(*38–40*). To enable facile fusion with diverse vaccine antigens and maximize co-receptor crosslinking, we engineered the monomeric CR2 binder CB7 into a trimeric assembly (Fig. 1E).

We retained the CR2-binding helical motif from CB7 and generated trimeric backbones under strict C3 symmetry, with three copies of the binding motif arranged along the z-axis. To avoid conformational constraints that would compromise core packing, we introduced rigid-body perturbation during design: we maintained C3 symmetry while sampling limited translational and rotational degrees of freedom of the binding motifs, to identify conformations that form a stable hydrophobic core while preserving the geometry required for CR2 recognition (Fig. S2A). Using the same design pipeline as for monomeric binders, we generated 4,000 trimeric CR2 binder designs that stably display the CR2-binding helices. From the 12 designs with highest structural convergence, we identified two candidates with strong CR2 binding and high protein stability, designated CB7t3 and CB7t4 (Fig. S2B), and BLI profiles of all 12 trimeric candidates are provided in Fig. S5.

We next characterized the biophysical properties of CB7, CB7t3, and CB7t4. Monomeric CB7 eluted as a single symmetric peak on size-exclusion chromatography (SEC), indicating high monodispersity. BLI measurements revealed a CR2 binding affinity (*K*_d_) of ∼26 nM. Thermal shift assays showed that CB7 retains its folded conformation up to ∼50 °C, demonstrating good structural stability (Fig. 1F).

The trimeric variants CB7t3 and CB7t4 also eluted as single monodisperse peaks on SEC, with elution volumes consistent with their larger trimeric molecular weight relative to CB7. BLI assays confirmed that both trimeric binders bound CR2 with ∼20-fold higher avidity (apparent *K*_d_) than monomeric CB7, consistent with multivalent engagement. Thermal stability was also improved, with unfolding onset above 65 °C for both trimers (Fig. 1G). Both CB7t3 and CB7t4 cross-reacted with murine CR2, consistent with the conserved SCR1-2 architecture across species (Fig. S3); apparent affinities toward mouse CR2 were in the low-nanomolar range (Fig. S4), supporting their use in preclinical mouse immunization studies.

### CR2 binders specifically engage B cells and enhance antigen-driven activation

To confirm that the designed binders recognize native CR2 on the B cell surface, we performed flow cytometry on two CR2-high B cell lines, Akata and Raji, using native human C3d (hC3d) as a positive control and the trimeric scaffold I53-50A1 (50A, a non-binding nanoparticle component) as a negative control(*41*). Both CB7t3 and CB7t4 exhibited significantly higher mean fluorescence intensity (MFI) than hC3d, while 50A showed no specific binding (Fig. 2A and 2B; Fig. S8B). Specificity for CR2 was further validated in HEK293T cells ectopically expressing human or murine CR2: CB7t3 and CB7t4 bound efficiently to both transfectants, while 50A remained at background levels (Fig. S7A to S7C). Confocal microscopy in Raji cells and freshly isolated primary human CD19^+^ B cells confirmed strong colocalization of SA-AF488-labeled CB7t3 with CD21/CR2 at the plasma membrane (Fig. 2C), further validating recognition of native cell-surface CR2.

**Fig. 2.**
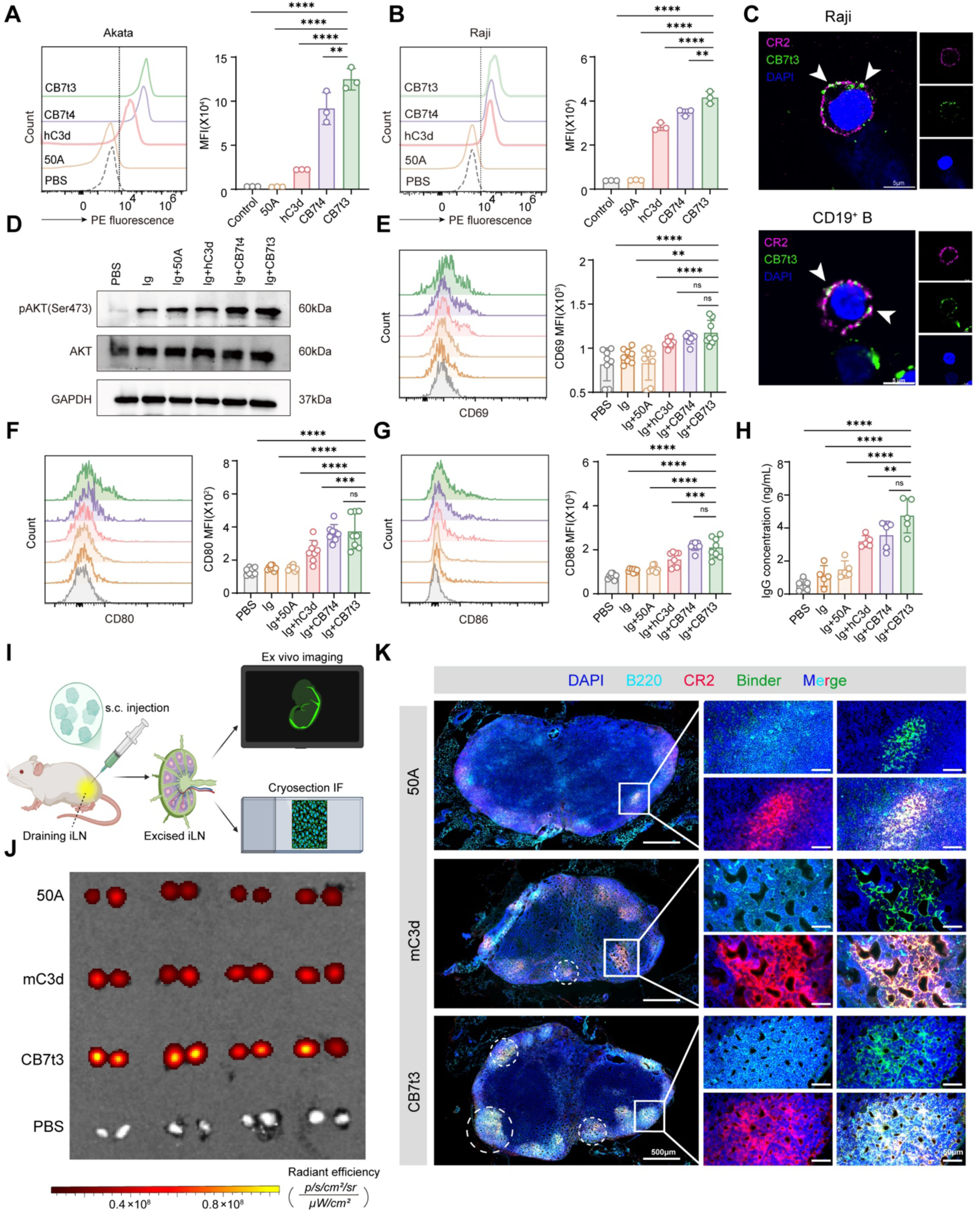
Designed CR2 binders amplify subthreshold BCR signaling and promote follicular localization of the scaffold. (A and B) Representative flow cytometry histograms and quantified mean fluorescence intensity (MFI) of PE-labeled CB7t3, CB7t4, hC3d, and 50A on Akata (A) and Raji (B) B cell lines expressing endogenous CR2; PBS served as the background control. (C) Confocal microscopy images of Raji cells and freshly isolated primary human CD19^+^ B cells stained with CD21-AF700 to detect endogenous CD21/CR2 (pseudocolored magenta), AF488-CB7t3 (green), and DAPI (blue). Arrowheads indicate colocalization of AF488-CB7t3 with CD21/CR2 at the plasma membrane. Scale bars are indicated. (D) Immunoblot analysis of phospho-AKT (Ser473) in primary human CD19^+^ B cells stimulated for 2 min with a suboptimal dose of anti-Ig alone or together with 50A, hC3d, CB7t4, or CB7t3. Total AKT and GAPDH are shown as controls. (E to G) Representative histograms and quantified MFI of the activation markers CD69 (E), CD80 (F), and CD86 (G) on primary human CD19^+^ B cells 48 h after stimulation with low-dose anti-Ig alone or together with the indicated proteins. (H) Total IgG concentrations in culture supernatants after 7 days of primary human CD19^+^ B cell stimulation under the indicated conditions. (I) Experimental scheme for subcutaneous administration of mCherry-fused proteins, followed by excision of draining inguinal lymph nodes 4 h later for *ex vivo* fluorescence imaging and cryosection immunofluorescence. (J) *Ex vivo* mCherry fluorescence images of draining inguinal lymph nodes from mice injected with mCherry-fused 50A, murine C3d (mC3d), or CB7t3, or with PBS; corresponding quantification is shown in Fig. S7E. Four mice were analyzed per group (*n* = 4), with bilateral inguinal lymph nodes collected from each mouse. (K) Representative multicolor immunofluorescence images of draining lymph node cryosections showing mCherry-fused 50A, mC3d, or CB7t3 (pseudocolored green), B220^+^ B cell follicles (cyan), CD21/CD35 (CR2/CR1; red), and DAPI-stained nuclei (blue). CB7t3 showed prominent enrichment within B220^+^ follicles and overlap with CD21/CD35-rich regions, whereas 50A and mC3d lacked comparable follicular targeting. Images are representative of four mice per group. Scale bars are indicated. Data in (A) and (B) are mean ± s.e.m. from three independent experiments (*n* = 3); data in (E) to (G) are mean ± s.e.m. from eight independent experiments (*n* = 8); and data in (H) are mean ± s.e.m. from five independent experiments (*n* = 5). Each symbol represents one independent experiment. Statistical significance was determined by one-way ANOVA with Tukey’s multiple-comparisons test. \**P* < 0.05; \*\**P* < 0.01; \*\*\**P* < 0.001; \*\*\*\**P* < 0.0001; ns, not significant.

We next tested whether CR2 engagement by the designed binders enhances BCR signaling under subthreshold activation conditions. Co-engagement of the CR2-CD19-CD81 co-receptor complex with BCR promotes phosphorylation of CD19 cytoplasmic tyrosine residues, recruits PI3K, and increases membrane PIP3 levels, driving AKT membrane recruitment and phosphorylation to amplify BCR signaling(*15–18*, *21*, *42*, *43*). We stimulated primary human CD19^+^ B cells with a low dose of BCR crosslinker together with CB7t3 or CB7t4. Both trimeric binders enhanced AKT Ser473 phosphorylation more strongly than hC3d (Fig. 2D).

Flow cytometric analysis showed that CB7t3 and CB7t4 upregulated the early activation marker CD69, as well as co-stimulatory molecules CD80 and CD86, to levels exceeding those induced by hC3d (Fig. 2E to 2G and Fig. S8C), consistent with enhanced B cell activation. CB7t3 and CB7t4 also increased total IgG secretion from cultured primary B cells and expanded the CD27^+^IgG^+^ memory-phenotype population relative to hC3d (Fig. 2H, Fig. S7D, and Fig. S8A), indicating enhanced IgG secretion and enrichment of a class-switched CD27^+^ phenotype relative to native C3d.

We next evaluated *in vivo* targeting efficacy (Fig. 2I), focusing on CB7t3, which showed superior *in vitro* performance. mCherry-fused CB7t3 accumulated rapidly in draining lymph nodes within 4 h after subcutaneous injection, at significantly higher levels than the corresponding 50A or murine C3d (mC3d) fusion controls (Fig. 2J and Fig. S7E). Multicolor immunofluorescence of draining lymph node sections revealed that mCherry-fused CB7t3 was enriched within B220^+^ follicles and showed substantial overlap with CD21/CD35-rich structures, whereas the 50A and mC3d fusion controls lacked comparable follicular targeting (Fig. 2K). These *in vivo* results confirm that CB7t3 efficiently targets B cells in lymphoid tissues, establishing its immunological suitability as a vaccine scaffold.

### Structural basis for CR2 recognition and trimer assembly

To validate the structural accuracy of our de novo design and define the molecular basis of CR2 binding, we determined the cryo-electron microscopy (Cryo-EM) structure of the CB7t3–CR2 (SCR1-2) complex at 2.97 Å resolution (Fig. S9, Table S1). The density map enabled unambiguous tracing of the polypeptide backbone and side chains, revealing a 3:3 binding stoichiometry where each CB7t3 protomer engages one CR2 molecule (Fig. 3A and Fig. S10). All-atom alignment of the experimental structure with computational design model validated by AlphaFold3 (AF3) yielded a root-mean-square deviation (RMSD) of only 0.427 Å, with near-identical side-chain conformations at the binding interface (Fig. 3B, 3D, and 3F; Fig. S12), confirming the near-atomic accuracy of our de novo design.

**Fig. 3.**
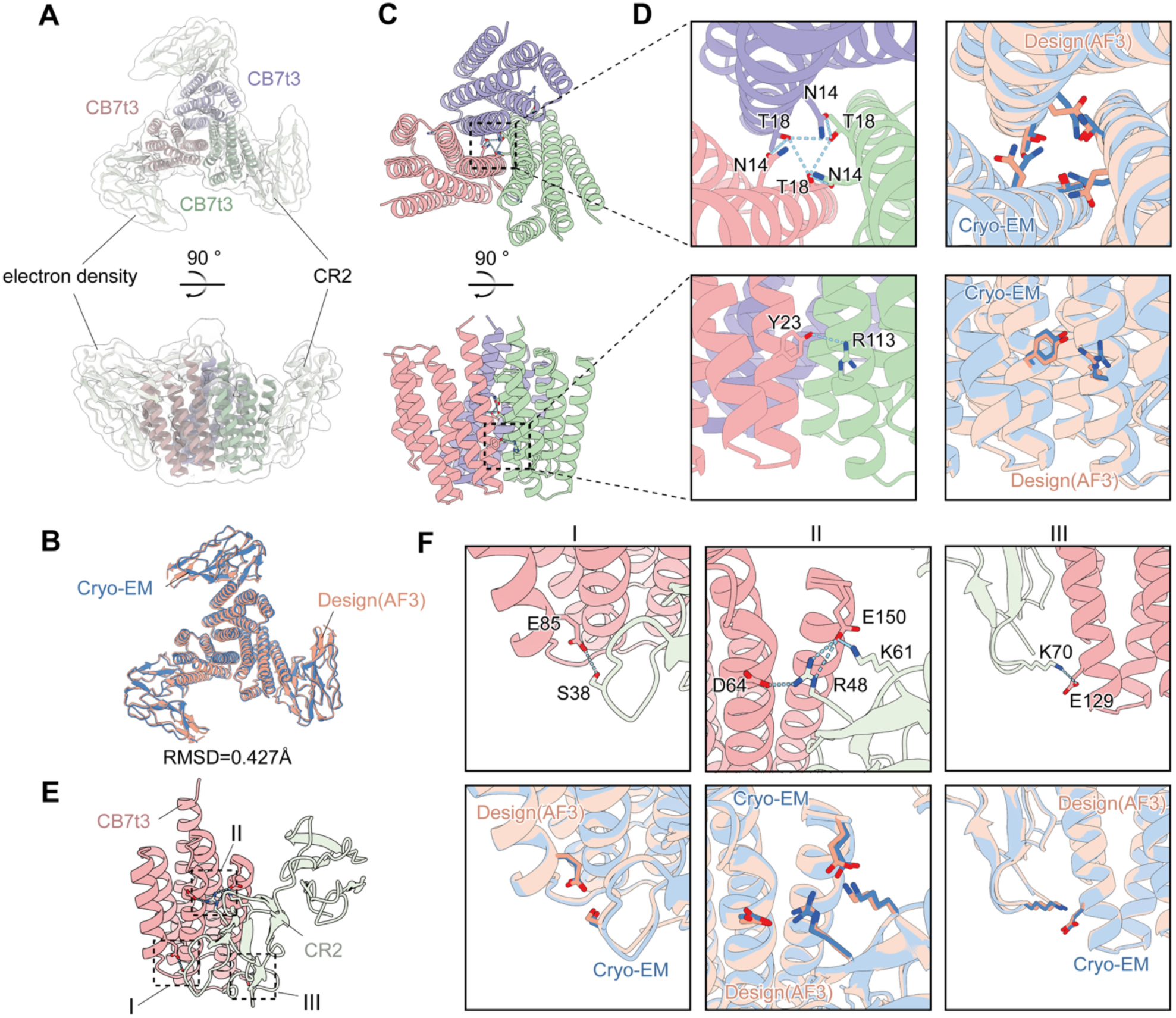
Cryo-EM structure of the CB7t3–CR2 complex reveals the molecular basis of trimer assembly and CR2 recognition. (A) Cryo-EM density map of the CB7t3–CR2 complex determined at 2.97 Å resolution. The central colored density corresponds to the rigid helical-bundle core of the CB7t3 trimer, and peripheral light-gray densities correspond to three radially arranged CR2 SCR1-2. This architecture minimizes steric interference between adjacent receptors and provides a structural basis for efficient B cell receptor co-crosslinking. (B) All-atom superposition of the experimentally determined cryo-EM structure and the AlphaFold3-validated design model. (C) Top and side views of the CB7t3 trimeric scaffold. Individual protomers are colored separately to illustrate the compact helical-bundle architecture. (D) Structural basis of trimer stabilization in CB7t3. Top left: the central hydrogen-bond network defining the trimeric assembly, in which Asn14 and Thr18 from the three protomers form a triangular polar lock. Bottom left: polar interactions at the lateral helical-bundle interface, where Tyr23 forms interchain hydrogen bonds with Arg113 from the adjacent protomer. Right: superposition of the designed model and the experimentally determined structure for key interface residues. (E) Close-up view of the CB7t3–CR2 binding interface, illustrating the compact and focused interaction mode. (F) Molecular interactions between CB7t3 and CR2, highlighting three major interaction clusters. Bottom: superposition of the designed model and the experimentally determined structure for key interface residues.

Analysis of the CB7t3 trimer core revealed extensive hydrophobic packing along the helical bundle (Fig. 3C). A central hydrogen-bond network, characterized by reciprocal N14–T18 interactions across the C3 symmetry axis, forms a triangular polar lock that minimizes assembly free energy and enforces strict trimer stoichiometry, thereby preventing dimerization or non-specific aggregation (Fig. 3C). This internal architecture is further reinforced by lateral hydrophobic packing, notably via a Y23–R113 inter-subunit hydrogen bond that serves as a critical binding hotspot. The congruence between the designed and experimental structures confirms that the trimer assembles exactly as modeled.

At the binding interface, CB7t3 engages CR2 through a multivalent network of polar interactions clustered into three distinct regions (Fig. 3E). Region I is anchored by a hydrogen bond between E85 (CB7t3) and S38 (CR2). Region II constitutes the major electrostatic anchoring site, comprising a D64-R48 salt bridge and a bifurcated interaction in which E150 (CB7t3) bridges R48 and K61 of CR2 (Fig. 3F and Fig. S11). Region III features a bidentate salt bridge between E129 (CB7t3) and K70 (CR2) (Fig. 3F). This multi-point polar interaction network is consistent with the *in vitro* binding data and rationalizes the high affinity and specificity of CB7t3 for CR2. Collectively, the agreement between designed and experimental binding interfaces supports the use of CB7t3 as a scaffold for vaccine engineering.

### The CR2 binder vaccine scaffold enhances humoral immunity

Epstein-Barr virus (EBV) remains without a licensed vaccine, due in part to the low immunogenicity of its envelope glycoproteins(*44–46*). We previously demonstrated that nanoparticle display substantially enhances the immunogenicity of EBV antigens, providing a benchmark for evaluating our CR2 binder vaccine scaffold(*45–48*). We selected the monomeric EBV glycoprotein complex gHgL as the model antigen, rather than the trimeric gB fusion protein(*49–51*), to avoid confounding effects from intrinsic antigen trimerization.

We expressed and purified five vaccine constructs: gHgL monomer, gHgL fused to the I53-50A1 trimeric scaffold (gHgL-50A), gHgL displayed on I53-50 icosahedral nanoparticles (gHgL-NP), gHgL fused to monomeric murine C3d (gHgL-mC3d), and gHgL fused to trimeric CB7t3 (gHgL-CB7t3) (fusion designs, SEC profiles, and SDS–PAGE analysis are shown in Fig. S6). Mice were immunized subcutaneously three times at 3-week intervals, with serum and tissue collected at week 8 (Fig. 4A).

**Fig. 4.**
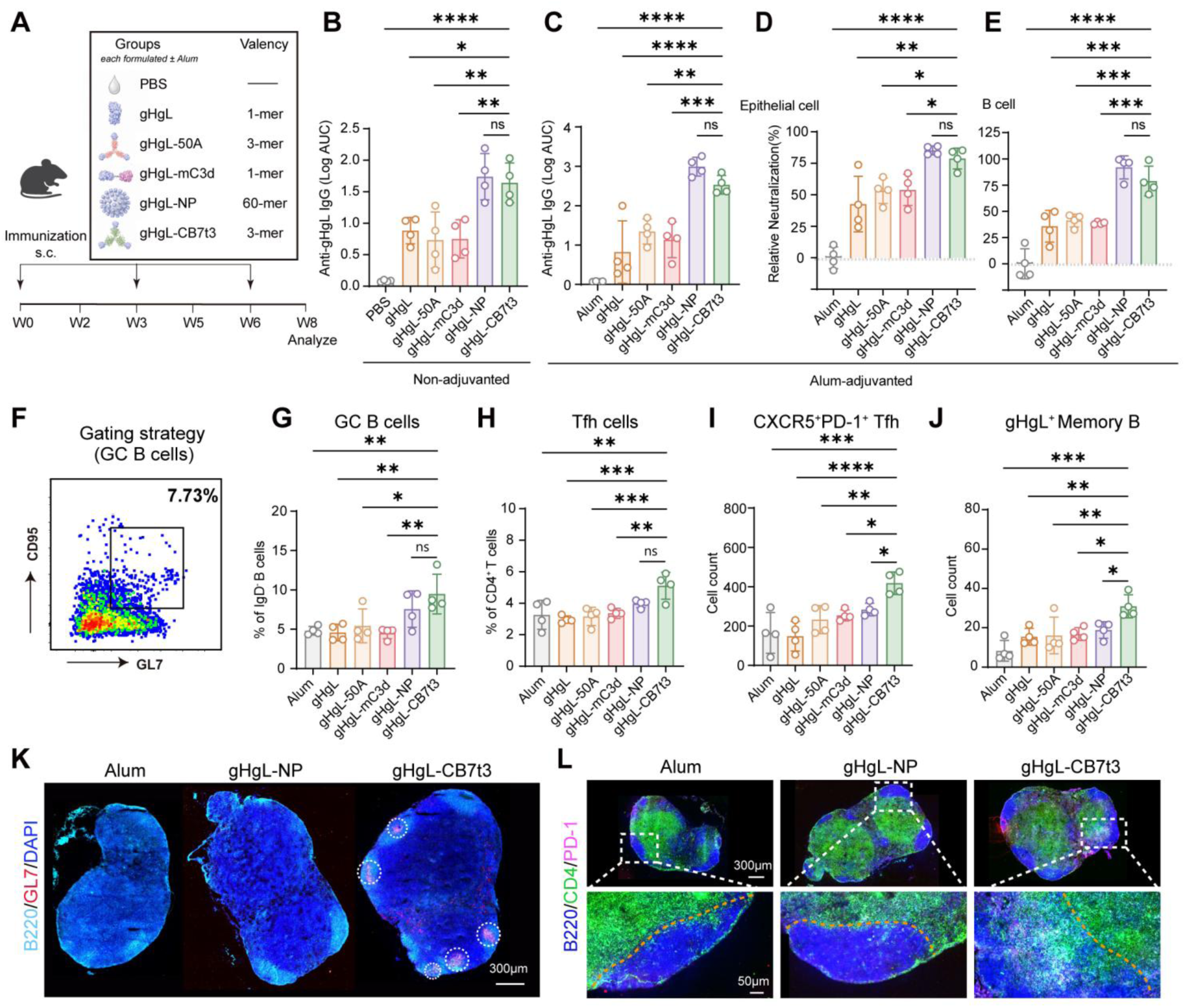
CR2 binder as vaccine scaffold elicits potent humoral immunity with enhanced germinal center and antigen-binding IgG^+^ B-cell responses. (A) Immunization scheme and vaccine groups. BALB/c mice were immunized subcutaneously at weeks 0, 3, and 6 in two parallel cohorts: a non-adjuvanted cohort receiving PBS or the indicated gHgL formulations in PBS, and an alum-adjuvanted cohort receiving alum alone or the indicated gHgL formulations formulated with alum. All immunogens were administered at an equal gHgL-molar dose. Serum was collected at weeks 2, 5, and 8, and immune responses were analyzed at week 8. Antigen valency per assembly is indicated for each formulation. (B and C) Week 8 serum anti-gHgL IgG responses, expressed as log-transformed area under the ELISA binding curve (Log AUC), in mice immunized without adjuvant (B) or with alum (C). (D and E) Relative neutralization (%) of EBV infection in epithelial cells (D) and B cells (E) by sera from alum-adjuvanted mice at the initial 1:100 serum dilution, calculated as 100 minus relative infection. (F) Representative terminal GL7-versus-CD95 plot gated on splenic B220^+^IgD^−^ B cells. (G to J) Flow-cytometric quantification after alum-adjuvanted immunization: (G) frequency of splenic GC B cells among IgD^−^ B cells; (H) frequency of splenic Tfh cells among CD4^+^ T cells; (I) splenic PD-1^+^ Tfh cell count; and (J) peripheral-blood gHgL-binding IgG^+^ memory-phenotype B cell count. For (I), equal numbers of input splenocytes were stained and equal total cellular events were acquired. For (J), equal numbers of input peripheral-blood lymphocytes were stained and equal total cellular events were acquired. Counts indicate cells within the terminal gates under standardized acquisition conditions. Representative splenic gating strategies for the corresponding GC B-cell, Tfh-cell, and gHgL-binding IgG^+^ B-cell analyses are shown in Fig. S15A, Fig. S16A, and Fig. S17A, respectively. (K) Representative draining lymph-node cryosections stained for B220 (cyan), GL7 (red), and DAPI (blue), showing GC structures after immunization with alum alone, gHgL-NP, or gHgL-CB7t3. Scale bar, 300 μm. These selected groups represent the adjuvant control and the two highest-performing vaccine formulations. (L) Representative whole-section and magnified images stained for B220 (cyan), CD4 (green), PD-1 (magenta), and DAPI (blue); boxed regions are magnified below. Scale bars, 300 μm (top) and 50 μm (bottom). Data are mean ± s.e.m.; n = 4 mice per group, and each symbol represents one mouse. Statistical significance was determined by one-way ANOVA with Tukey’s multiple-comparisons test. \**P* < 0.05; \*\**P* < 0.01; \*\*\**P* < 0.001; \*\*\*\**P* < 0.0001; ns, not significant.

In the absence of exogenous adjuvant, gHgL-CB7t3 and gHgL-NP induced the highest anti-gHgL IgG responses (Log AUC) at week 8, significantly exceeding those of gHgL-50A (non-targeted trimer) and gHgL-mC3d (targeted monomer), with no body weight changes across groups (Fig. 4B and Fig. S13A). Splenic germinal center (GC) B cell and follicular helper T (Tfh) cell frequencies did not differ significantly among groups, consistent with the weak immunogenicity of non-adjuvanted protein antigens(*6*) (Fig. S13B and S13C). Splenic memory B cell and T cell memory subsets showed upward trends in the gHgL-CB7t3 and gHgL-NP groups, although most comparisons did not reach statistical significance (Fig. S13D and S13E; Fig. S14A to S14D). These data demonstrate that CR2 targeting enhances antibody responses to protein antigens in the absence of adjuvant, with efficacy comparable to nanoparticle display.

We next tested the vaccine constructs formulated with aluminum hydroxide (alum) adjuvant, to evaluate compatibility with clinically established adjuvant systems (Fig. 4A). Alum boosted gHgL IgG responses across all groups; gHgL-CB7t3 and gHgL-NP remained the top performers, with antibody levels far exceeding those of gHgL-50A and gHgL-mC3d (Fig. 4C). EBV neutralization assays showed that sera from the gHgL-CB7t3 and gHgL-NP groups produced similarly high relative neutralization of both epithelial-cell and B-cell infection, outperforming the other groups (Fig. 4D and 4E). Thus, fusion to the CR2 binder vaccine scaffold markedly increases antigen immunogenicity. Under the equal gHgL-molar dosing used here, the trimeric CR2 binder achieved responses comparable to the icosahedral nanoparticle formulation, while outperforming both simple trimerization and native C3d fusion strategies.

To dissect the immunological features associated with the enhanced humoral response, we analyzed immune cell populations in alum-immunized mice. gHgL-CB7t3 induced significantly higher frequencies of splenic GC B cells than all other groups except the gHgL-NP group (Fig. 4F and 4G; Fig. S15A and S15B). It also increased splenic Tfh cell frequencies, and the number of acquired PD-1^+^ Tfh events under standardized flow-cytometric acquisition was higher than that in the gHgL-NP group (Fig. 4H and 4I; Fig. S16A to S16C), consistent with an enhanced GC-associated response. In peripheral blood, gHgL-CB7t3 also increased the number of acquired gHgL-binding IgG^+^ memory-phenotype B-cell events relative to gHgL-NP under standardized acquisition conditions (Fig. 4J). Consistent with this finding, splenic analysis also showed an increased gHgL-binding IgG^+^ memory-phenotype B-cell population in the gHgL-CB7t3 group (Fig. S17A to S17C).

These flow-cytometric findings were supported by immunofluorescence analysis of draining lymph nodes. gHgL-CB7t3 immunization produced more prominent GL7^+^ GC structures and increased CD4^+^PD-1^+^ signal within B220^+^ follicular regions (Fig. 4K and 4L; Fig. S16D and S16E). Together, these results indicate that gHgL-CB7t3 is associated with enhanced GC B-cell and Tfh responses in both the spleen and draining lymph nodes. Beyond these GC-associated changes, splenic analysis revealed increases in CD4^+^ effector and central memory T-cell populations, as well as CD8^+^ effector memory T-cell populations, following gHgL-CB7t3 immunization (Fig. S18A to S18D), suggesting that the vaccine was also accompanied by broader changes in the splenic T-cell compartment. To further assess whether immunization with the CB7t3 scaffold induces cross-reactive antibodies against host complement components, we quantified anti-mC3d IgG responses in immune sera. In both non-adjuvanted and alum-adjuvanted cohorts, the gHgL-mC3d group elicited substantially elevated anti-mC3d IgG responses, with both binding curves and AUC values exceeding those of all other vaccine groups. In contrast, anti-mC3d IgG levels in the gHgL-CB7t3 group showed no significant increase and were overall comparable to those in control groups including gHgL, gHgL-50A, and gHgL-NP (Fig. S19A to S19D). We next performed *ex vivo* fluorescence imaging of draining lymph nodes and major organs after subcutaneous injection of mCherry-fused CB7t3. The CB7t3-mCherry signal was predominantly enriched in the draining inguinal lymph nodes, with markedly lower fluorescence detected in other major organs (Fig. S20). These results demonstrate that CB7t3 does not induce detectable C3d cross-reactivity and exhibits a tissue-distribution profile characterized by preferential accumulation in draining lymph nodes and low exposure to systemic organs. These data provide preliminary support for the safety of the CR2-targeting vaccine scaffold at both the immunological cross-reactivity and *in vivo* biodistribution levels and highlight a potential advantage of the de novo-designed CR2-binding scaffold over direct native mC3d fusion in limiting anti-complement antibody responses.

Collectively, the CR2 binder vaccine scaffold achieves humoral immunity comparable to state-of-the-art nanoparticle platforms, while operating through a distinct immunological mechanism that is associated with enhanced GC-associated responses and increased antigen-binding IgG^+^ B-cell events. The complementary modes of action leave open the possibility of combining CR2 targeting with nanoparticle display for even greater vaccine efficacy.

## Discussion

The CR2-CD19 co-receptor complex is well established as a central regulator of the B cell activation threshold, and its engagement represents one of the most potent mechanisms to amplify antigen-specific humoral responses(*15*, *16*, *19*, *21*). Despite this well-established biological rationale, clinical translation of CR2-targeted vaccine strategies has remained stalled for decades, constrained by the inherent limitations of native C3d as a targeting ligand(*23*, *24*, *31*). Monomeric C3d binds CR2 with only micromolar affinity, requiring tandem multimerization to achieve meaningful adjuvant effects(*13*, *52*, *53*); it also suffers from poor biophysical stability, challenges in large-scale manufacturing, and inherent risks of disrupting endogenous complement homeostasis or triggering autoreactive responses(*13*, *25*, *54*). Alternative B cell–targeting approaches, such as anti-CD19 or anti-BCR antibody conjugates, add substantial manufacturing complexity and cost, and often fail to recapitulate the physiological co-stimulatory signaling of natural complement opsonization(*16*, *18*). To date, no CR2-targeting modality has combined high affinity, favorable developability, and safety into a platform suitable for broad vaccine application, leaving a major axis of B cell regulation untapped for rational vaccine engineering.

Our CR2 binder vaccine scaffold directly fills this technological void. Built on a de novo protein framework with no sequence homology to endogenous complement components, it achieves nanomolar CR2 binding affinity without competing with physiological complement function at relevant doses, mitigating the safety liabilities of native C3d-based strategies. When formatted as a trimer, the scaffold achieves multivalent co-receptor crosslinking that mimics clustered complement-opsonized antigen, delivering potent co-stimulatory signals in parallel with BCR engagement. In head-to-head comparison with a native C3d fusion vaccine, the trimeric CR2 binder induced substantially higher B cell activation, germinal center formation, and antigen-specific antibody production, demonstrating that engineered binders can outperform natural complement fragments as vaccine adjuvant modules. Notably, the scaffold’s ability to actively home to B cell follicles in draining lymph nodes adds a second layer of enhancement beyond co-stimulation, concentrating antigen at the site of B cell priming to further improve response magnitude. Cryo-EM determination of the CB7t3–CR2 complex at 2.97 Å resolution confirmed that the experimental structure matches the AlphaFold3-validated design model to within 0.43 Å RMSD, supporting the fidelity of the de novo design pipeline and the predictability of CR2-binding geometry.

When evaluated against self-assembling protein nanoparticles, the current trend for subunit vaccine multimerization, the CR2 binder scaffold achieves comparable relative neutralization levels through a fundamentally different mechanism(*55*, *56*). Whereas nanoparticles passively concentrate BCR crosslinking by antigen density, the binder scaffold actively directs antigen to CR2^+^ B cells and engages the co-receptor complex in a defined geometry. The two modes of action are complementary rather than competitive: the binder can be displayed on nanoparticle surfaces to combine active B cell targeting with high-density antigen presentation. Beyond this conceptual distinction, the scaffold offers practical translational advantages. It is produced as a single polypeptide chain compatible with standard recombinant expression and purification workflows, exhibits high thermal stability that simplifies formulation and storage, and accommodates monomeric or trimeric antigens via flexible genetic fusion without re-engineering the scaffold core (Fig. S6). Together, these properties create a clear upgrade path from early-stage candidates to maximally potent clinical formulations.

Beyond prophylactic antiviral vaccines, this design paradigm, engineering de novo binders to engage immune co-receptors in a defined multivalent format, can in principle be extended to other stimulatory pathways on B cells or other immune cell types. Whether such extensions translate to therapeutic settings such as chronic infection or cancer immunotherapy will require dedicated disease models and was not tested in the present work. Because the SCR1-2 ectodomains of human, mouse, and bovine CR2 are structurally conserved (Fig. S3), the same design principle may extend to veterinary vaccines, although species-specific validation will be required. Moreover, the current trimeric configuration can be extended through additional oligomerization modules to generate higher-order assemblies with increased valency, allowing the immune activation strength to be tuned for different immunological contexts.

Further work is required to test the scaffold across diverse antigens and species, to evaluate long-term safety and anti-binder antibody responses after repeated dosing, particularly given that any de novo protein scaffold could in principle elicit anti-binder antibodies that limit efficacy upon boosting(*57*), and to assess compatibility with clinically approved adjuvants beyond alum. Although our three-dose regimen did not show overt suppression of antigen-specific antibody responses, dedicated assays for anti-CB7t3 IgG titers and cross-reactivity to the host proteome will be required before clinical translation. The combination of single-chain manufacturability, structural tunability, and CR2-specific co-stimulation positions this platform as a practical starting point for next-generation protein vaccines.

Taken together, this work establishes CR2 as a druggable node for rational vaccine engineering and demonstrates that de novo protein design can deliver what three decades of complement-based approaches could not: a high-affinity, manufacturable, and structurally defined co-receptor agonist. By coupling antigen display to precise co-receptor engagement within a single polypeptide, the CR2 binder scaffold converts a fundamental principle of B cell biology into a practical engineering module. More broadly, the design strategy presented here—atomic-level mimicry of a natural immune interaction with a fully synthetic protein—provides a general blueprint for harnessing immune co-receptors, and brings mechanism-guided vaccine design a step closer to clinical reality.

## Materials and Methods

### Monomeric binder design

The CR2–C3d interface in the crystal structure (PDB: 3OED) was analyzed using the Rosetta InterfaceAnalyzer protocol(*34*), followed by hotspot residue identification using PDBePISA(*35*). Based on the physicochemical properties of interface residues and the local structural environment surrounding CR2–C3d hotspot residues, residues suitable for guiding binder generation were selected. Five hotspot residues, including L57, L64, I66, Y36, and V46, were defined as spatial constraints for binder design.

RFdiffusion(*26*) was used for de novo backbone generation using the pretrained RFdiffusion model weights. A total of 12,000 candidate binder backbones were generated with complete diffusion trajectories consisting of 50 denoising steps. The designed binders were constrained to a sequence length range of 120–160 amino acids. The generated backbones were filtered based on structural integrity, interface geometry, and spatial arrangement. Designs containing fewer than four α-helices were removed. ProteinMPNN(*27*, *58*) was subsequently used to design amino acid sequences for the filtered backbones using pretrained soluble-protein model weights. For each backbone, four corresponding sequences were generated. The resulting backbone–sequence pairs were evaluated using Chai-1(*36*) for structural fidelity and target-binding compatibility. CR2–binder complexes were directly predicted without multiple sequence alignment (MSA) searching. Designs satisfying the following criteria were retained: interface predicted template modeling score (ipTM) > 0.80, predicted template modeling score (pTM) > 0.85, and backbone RMSD between the designed and predicted structures below 2.0 Å.

The retained binder sequences were subsequently evaluated using AlphaFold3(*59*) complex prediction with MSA searching enabled. Models with ipTM > 0.80 were subjected to interface analysis using PDBePISA(*35*). Designs exhibiting calculated interface ΔG contributions greater than 8 kcal/mol were retained. Candidate binders were ranked according to structural agreement, interface geometry, and Rosetta energy scores, and the top five designs were selected for further optimization.

### Monomeric binder optimization

The five highest-ranked parental binders were subjected to partial-diffusion refinement. For each parental design, 4,000 optimized backbone models were generated using RFdiffusion. The sequence length was maintained identical to the parental design. Partial diffusion was performed using 15 noise steps followed by 15 denoising steps. The refined backbones were filtered using criteria similar to those applied during the initial design campaign. ProteinMPNN was subsequently used for sequence redesign of the filtered backbones using pretrained soluble-protein model weights, generating two sequences per backbone.

The optimized backbone–sequence pairs were evaluated using the same structure–sequence screening pipeline with more stringent thresholds. Designs were required to achieve ipTM > 0.85 and backbone RMSD below 1 Å. AlphaFold3-based complex predictions were further filtered using an ipTM threshold of >0.85. Following computational ranking, the top 12 optimized designs were selected for experimental characterization. Three designs exhibited strong binding activity toward both human and mouse CR2. Among these candidates, CB7 showed the most favorable combination of binding affinity and protein solubility.

### C3-symmetric trimer design

The cross-reactive monomeric binders identified above were used as templates for the design of C3-symmetric trimeric binders through a motif-scaffolding strategy. The CR2-binding motifs were arranged around the C3 symmetry axis (z-axis), and four degrees of freedom were randomly sampled to explore possible spatial configurations. Configurations with average inter-protomer distances ranging from 16 to 28 Å were selected as reference templates.

RFdiffusion was subsequently used to generate trimeric scaffold backbones based on these reference arrangements using the pretrained RFdiffusion base model. The intra-chain and inter-chain interaction weights were set to weight_intra = 1.0 and weight_inter = 0.06, respectively. A total of 4,000 trimeric scaffold backbones were generated with sequence lengths ranging from 123 to 158 amino acids. ProteinMPNN was directly applied for sequence design of the generated homotrimeric assemblies, generating two sequences per backbone.

The resulting six-chain complexes (3×CR2–3×binder) were evaluated using Chai-1 without MSA searching. Designs satisfying the following criteria were retained: ipTM > 0.85, pTM > 0.85, and backbone RMSD between designed and predicted structures below 1.5 Å. The retained trimeric binder sequences were further evaluated using AlphaFold3 complex prediction with MSA searching enabled. Models with ipTM > 0.85 were analyzed using PDBePISA, and designs exhibiting calculated trimer interface ΔG contributions greater than 10 kcal/mol were retained.

Final candidates were ranked based on structural agreement, interface geometry, and Rosetta energy scores. The top 12 designs were selected for experimental characterization. Two designed trimeric proteins exhibited high structural stability and high-affinity binding toward both human and mouse CR2.

### Cell lines and culture conditions

Akata and Raji B cell lines were cultured in RPMI 1640 medium (Gibco, cat. no. C11875500BT) supplemented with 10% (v/v) fetal bovine serum (FBS; ExCell Bio, cat. no. FSP500). HEK293 and HEK293T cells were maintained in high-glucose Dulbecco’s modified Eagle’s medium (DMEM; Gibco, cat. no. C11995500BT) supplemented with 10% FBS. Unless otherwise indicated, media contained 1% (v/v) penicillin–streptomycin (Gibco, cat. no. 15140-122), and cells were maintained at 37 °C in 5% CO₂.

HEK293F cells were cultured in serum-free FreeStyle 293F expression medium (Sino Biological, cat. no. M293TII-1) in baffled Erlenmeyer flasks at 37 °C, 5% CO₂, and 120 rpm. Cell lines were passaged within the recommended density range and routinely tested for mycoplasma contamination by PCR.

### Human blood samples and isolation of primary B cells

Peripheral blood was obtained from healthy adult donors after written informed consent under a protocol approved by the institutional ethics committee of Sun Yat-sen University Cancer Center. Peripheral blood mononuclear cells were isolated by density-gradient centrifugation over Lymphoprep (Axis-Shield) at 400g for 20 min at room temperature. The mononuclear cell layer was collected and washed twice with PBS. CD19^+^ B cells were enriched by positive magnetic selection using the human CD19 MicroBead kit (Miltenyi Biotec, cat. no. 130-050-301) according to the manufacturer’s instructions. Purity was confirmed by flow cytometry and exceeded 90% before use. Independent donors were treated as biological replicates.

### Animals

Female BALB/c mice were purchased at 6 weeks of age from Beijing Vital River Laboratory Animal Technology. Mice were maintained under specific-pathogen-free conditions at the Sun Yat-sen University Cancer Center animal facility, acclimatized for at least 1 week, and randomly assigned to experimental groups before immunization or protein-administration studies. Randomization was performed using a computer-generated random sequence (Excel RAND function). Investigators conducting flow cytometry gating and lymph node immunofluorescence quantification were blinded to group assignment; blinding was lifted only after quantitative analysis was completed. All procedures were approved by the Institutional Animal Care and Use Committee of Sun Yat-sen University Cancer Center and were conducted in accordance with institutional and national guidelines. Group sizes were determined by power analysis (G*Power 3.1) to detect a 1.5-fold difference in Log AUC between groups with α = 0.05 and power = 0.80, informed by pilot variance from prior immunization studies. The minimal group size was *n* = 4 per arm.

### Expression plasmids and immunogen construction

Codon-optimized genes encoding the SCR1–2 ectodomains of human and mouse CR2 were cloned into mammalian expression vectors as SUMO-His fusion constructs for recombinant protein production. In parallel, SCR1–2 ectodomains were fused to the Fc region of immunoglobulin G for BLI-based binding assays. Genes encoding human C3d, mouse C3d, CB7, CB7t3, CB7t4, the nonbinding I53-50A1 trimeric scaffold (50A), and the I53-50B assembly component were cloned into bacterial expression vectors containing appropriate affinity tags for purification. His-tagged proteins were generated using vectors encoding a C-terminal polyhistidine (His₆) tag, whereas GST-fused proteins were constructed using the pGEX-6P-1 expression vector, which encodes an N-terminal GST tag followed by a PreScission protease cleavage site. For *in vivo* trafficking and biodistribution experiments, CB7t3, mouse C3d, and 50A were genetically fused in frame to mCherry.

The EBV gHgL antigen was derived from strain B95-8 and expressed from a bicistronic gH–furin cleavage site–P2A–gL cassette. The soluble gH and gL ectodomains were engineered for secretion, and the gL-containing product carried the purification tag. Mouse C3d, 50A, CB7t3, or CB7t4 was fused to the C terminus of gL through a 6xHis tag and a flexible G₄S linker to generate gHgL-mC3d, gHgL-50A, gHgL-CB7t3, and gHgL-CB7t4, respectively. gHgL-50A was combined with I53-50B to assemble the icosahedral gHgL nanoparticle (gHgL-NP). All expression cassettes were verified across the full insert by Sanger sequencing.

### Recombinant protein expression and purification

#### Mammalian protein expression and purification

Secreted proteins were produced by transient transfection of HEK293F suspension cells. Cells were maintained at a density of 1.0 × 10⁶ cells ml⁻¹ and transfected with plasmid DNA using linear polyethylenimine (PEI; Polysciences, cat. no. 40872) at a PEI:DNA mass ratio of 3:1 in Opti-MEM reduced-serum medium (Gibco, cat. no. 31985062). Cultures were incubated for 7 days at 37 °C with 5% CO₂ and shaking at 120 rpm. Supernatants were clarified by centrifugation (6,000 × g, 60 min, 4 °C) and filtered through a 0.45-μm membrane. His-tagged proteins were purified by Ni²⁺–NTA affinity chromatography (Cytiva, cat. no. 17525501), followed by size-exclusion chromatography (SEC) in PBS (pH 7.4). Protein purity was assessed by SDS–PAGE, and protein concentrations were determined from A₂₈₀ using sequence-derived molar extinction coefficients.

#### Bacterial expression and purification of His-tagged proteins

His-tagged proteins were expressed in *Escherichia coli* Rosetta cells. A single colony was inoculated into 10 ml of TB medium and cultured overnight. The starter culture was diluted 1:100 into fresh TB medium and grown at 37 °C with shaking at 110 rpm until an OD₆₀₀ of approximately 0.6–0.8 was reached. Cultures were cooled to 18 °C, and protein expression was induced with 1 mM IPTG. Cells were incubated at 18 °C with shaking at 110 rpm for approximately 16 h before harvesting by centrifugation (3,500 × g, 15 min, 4 °C).

Cell pellets were resuspended in TBS supplemented with protease inhibitor cocktail (25 ml per liter of culture) and lysed using TieChui™ E.coli Lysis Buffer (ACE Biosciences, BR0005-02). Lysates were incubated on ice for 15–30 min, clarified by centrifugation, and filtered through a 0.22-μm membrane. Clarified lysates were incubated with Ni²⁺–NTA affinity resin for 15 min at 4 °C. After washing with PBS containing 150 mM NaCl and 50 mM imidazole, bound proteins were eluted with PBS containing 150 mM NaCl and 500 mM imidazole. Eluted proteins were buffer-exchanged and concentrated by ultrafiltration.

#### Bacterial expression and purification of GST-fusion proteins

GST-fusion proteins were expressed in *Escherichia coli* Rosetta cells using the same expression protocol as described above. Briefly, cultures were grown in TB medium at 37 °C until an OD₆₀₀ of approximately 0.6–0.8 was reached, induced with 1 mM IPTG, and incubated at 18 °C with shaking at 110 rpm for approximately 16 h before harvesting by centrifugation (3,500 × g, 15 min, 4 °C).

Cell pellets were resuspended in TBS supplemented with protease inhibitor cocktail and lysed using TieChui™ E.coli Lysis Buffer (ACE Biosciences, BR0005-02). Lysates were clarified by centrifugation, filtered through a 0.22-μm membrane, and incubated with glutathione affinity resin for 1 h at 4 °C. After washing with TBS, GST-fusion proteins were eluted with 0.2 M reduced glutathione (GSH) prepared in TBS. Eluted proteins were buffer-exchanged and concentrated by ultrafiltration.

### Assembly and purification of gHgL nanoparticles

Purified gHgL-50A and I53-50B were combined at the validated I53-50 assembly stoichiometry and incubated under nondenaturing conditions to permit spontaneous formation of the 120-subunit I53-50 icosahedral assembly displaying 60 copies of gHgL. Assembled particles were separated from unassembled components and aggregates by preparative size-exclusion chromatography. Fractions corresponding to the principal nanoparticle peak were pooled, concentrated, and evaluated by analytical size-exclusion chromatography and SDS–PAGE before immunization. The gHgL-NP dose was adjusted according to the number of displayed gHgL copies and the molecular mass of the assembled immunogen to match the gHgL molar dose used for the other immunogens(*41*).

### SDS–PAGE and analytical size-exclusion chromatography

Purified proteins were mixed with 5× SDS sample buffer, heated at 98 °C for 10 min, and resolved on 10% polyacrylamide gels at 120 V for approximately 40 min together with a prestained molecular mass marker (Thermo Fisher Scientific, cat. no. 26616). Gels were stained with Coomassie Fast Blue solution (ACE Biosciences, cat. no. BR0070-01) for 30 min and destained in water. Images were acquired with a ChemiDoc MP system (Bio-Rad).

The oligomeric state and monodispersity of soluble proteins were assessed on Superdex 75 Increase 10/300 GL and Superdex 200 Increase 10/300 GL column connected to an ÄKTA Pure system (Cytiva). Larger assemblies were analyzed on a size-exclusion column with an exclusion range suitable for I53-50 nanoparticles. Chromatography was performed at 4 °C in PBS (pH 7.4) at 0.5 ml min⁻¹, and elution was monitored at 280 nm. Peak fractions were evaluated by their elution behavior and SDS–PAGE.

### Biolayer interferometry

Binding kinetics were measured on an Octet R8 instrument (Sartorius) at 30 °C with shaking at 1,000 rpm. PBS containing 0.1% (v/v) Tween-20 was used as kinetics buffer. Purified human or mouse CR2 SCR1–2 was immobilized on biosensors through its affinity tag to a loading response of approximately 0.6 nm. After a 30-s baseline, sensors were transferred to serial dilutions of binder for a 100-s association phase and then to kinetics buffer for a 100-s dissociation phase. First-round monomeric binders were screened at 5 μM. Trimeric candidates were characterized using seven twofold dilutions beginning at 400 nM; concentration ranges were adjusted when necessary to capture the full response window.

Signals from a ligand-loaded reference sensor exposed to buffer and from a nonbinding or analyte-only control were subtracted. Association and dissociation traces were globally fitted to a 1:1 binding model in Octet Analysis Studio. *K*_d_ values reported for trimeric binders represent apparent affinities that include avidity from multivalent CR2 engagement(*60*).

### Protein thermal stability

Thermal stability was measured on a UNcle instrument (Unchained Labs) by intrinsic differential scanning fluorimetry and static light scattering. Proteins were prepared in PBS at 1.0 mg ml⁻¹ and loaded into UniTube capillaries in 9-μl volumes. Samples were heated from 25 to 95 °C at 1 °C min⁻¹. Intrinsic tryptophan fluorescence was excited at 266 nm, and aggregation was monitored by static light scattering at 473 nm. Apparent melting temperatures were derived from the first derivative of the barycentric mean fluorescence signal, and aggregation onset was determined from the light-scattering trace. Measurements were performed in technical triplicate.

### Fluorescent labeling of recombinant proteins

Purified AviTag-containing CB7t3, CB7t4, 50A, human C3d (hC3d), and gHgL were site-specifically biotinylated with biotin-protein ligase BirA (GeneCopoeia, cat. no. BI001) according to the manufacturer’s instructions. Proteins were incubated with BirA in the supplied 1× Biotin Ligase Buffer A and 1× Biotin Ligase Buffer B containing ATP, magnesium acetate, and D-biotin at 30 °C for 30–40 min. Unincorporated biotin and other low-molecular-weight reaction components were removed by repeated dilution with PBS and concentration in centrifugal ultrafiltration devices. Biotinylated CB7t3, CB7t4, 50A, and hC3d were incubated with PE-conjugated streptavidin (SA-PE; 1 mg ml^−1^; Bioss, cat. no. bs-0437P-PE) protected from light. For comparisons, labeling reactions were normalized on a molar assembly basis (monomeric hC3d or trimeric CB7t3, CB7t4, and 50A), and a constant SA-PE-to-protein-assembly molar ratio and identical incubation conditions were used across preparations. The resulting complexes were designated PE-CB7t3, PE-CB7t4, PE-50A, and PE-hC3d, respectively, and were used for cell-surface binding assays. For confocal microscopy, biotinylated CB7t3 was separately complexed with streptavidin–Alexa Fluor 488 (SA-AF488) under light-protected conditions and was designated AF488-CB7t3. Biotinylated gHgL was separately incubated with streptavidin-conjugated BF750 (SA-BF750; Bioss, streptavidin cat. no. bs-0437P, BF750-conjugated format) protected from light, and the resulting BF750-gHgL probe was used for mouse lymphocyte staining.

### Binding to endogenous and ectopically expressed CR2

For analysis of endogenous CR2 binding, 1 × 10⁶ Akata or Raji cells were incubated on ice for 30 min with PE-labeled CB7t3, CB7t4, hC3d, or 50A. PBS-treated cells were used to define background. Cells were washed three times with cold PBS or FACS buffer and analyzed immediately by flow cytometry. Binding was quantified as mean fluorescence intensity among live singlets. Representative gating strategy is shown in Fig. S8B.

For cross-species binding, HEK293T cells were transfected with plasmids encoding full-length human or mouse CR2 together with an mCherry reporter. At 48 h after transfection, cells were detached, washed, and incubated on ice for 30 min with 1 μg of the indicated PE-labeled test protein per 1 × 10⁶ cells in 1 ml PBS. After three washes, binding was quantified as the percentage of live cells double-positive for the mCherry CR2 reporter and the PE-labeled test protein. Untransfected cells and cells incubated with PBS or 50A served as negative controls. Detailed gating strategy is shown in Fig. S7A.

### Confocal analysis of CR2 colocalization

Raji cells and freshly isolated primary human CD19^+^ B cells were allowed to settle on poly-L-lysine-coated glass-bottom dishes (NEST Biotechnology, cat. no. 801001). Cells were fixed with 4% paraformaldehyde for 10 min at room temperature, washed, and blocked with 5% BSA in PBS. Endogenous CR2 was detected with a directly conjugated anti-human CD21–Alexa Fluor 700 antibody (CD21-AF700; 1:50). AF488-CB7t3 was then added at 1 μg ml^−1^, and nuclei were counterstained with DAPI. Images were acquired on a Zeiss LSM 900 confocal microscope with a 63× oil-immersion objective using identical acquisition settings for samples compared within an experiment.

### Primary human B cell stimulation

Purified CD19^+^ B cells were cultured in 96-well U-bottom plates in complete RPMI 1640 medium at 1 × 10⁵ cells per well. BCR stimulation was initiated with 0.625 μg ml⁻¹ F(ab′)₂ goat anti-human IgG/IgM (H+L) (eBioscience, cat. no. 16-5099-85), either alone or together with 25 μg ml⁻¹ hC3d, CB7t3, CB7t4, or 50A. Unstimulated and anti-Ig-only controls were included in each experiment. Cells were stimulated for 2 min and immediately lysed for immunoblot analysis of AKT phosphorylation. Parallel cultures were harvested after 48 h for flow-cytometric analysis of activation markers and after 7 days for quantification of IgG secretion and analysis of CD27^+^IgG^+^ memory-phenotype B cells.

### Immunoblot analysis of AKT phosphorylation

Stimulation was terminated after 2 min by rapidly transferring the cells onto ice. Cells were lysed for 20 min in ice-cold RIPA buffer (Beyotime, cat. no. P0013B) supplemented with protease inhibitor cocktail (Roche, cat. no. 04693132001) and phosphatase inhibitor cocktail (Selleck, cat. no. B15002). Lysates were clarified by centrifugation at 12,000g for 15 min at 4 °C, and protein concentrations were determined using a BCA protein assay kit (Thermo Fisher Scientific, cat. no. 23225). Equal amounts of protein were resolved by SDS–PAGE and transferred to PVDF membranes (Millipore, cat. no. IPVH00010) at 250 mA for 90 min at 4 °C.

Membranes were blocked with 5% nonfat milk in Tris-buffered saline containing 0.1% Tween-20 (TBST) for 1 h at room temperature and incubated overnight at 4 °C with rabbit monoclonal antibody to phospho-AKT Ser473 (Cell Signaling Technology, cat. no. 4060T; 1:1,000) and mouse monoclonal antibody to GAPDH (clone 1E6D9; Proteintech, cat. no. 60004-1-Ig; 1:2,000). After washing with TBST, membranes were incubated for 1 h at room temperature with HRP-conjugated goat anti-rabbit IgG (H+L) (Proteintech, cat. no. SA00001-2; 1:5,000) and HRP-conjugated goat anti-mouse IgG (H+L) (Proteintech, cat. no. SA00001-1; 1:5,000).

After chemiluminescent detection of phospho-AKT and GAPDH, the same membranes were immersed in PBS containing Tween-20 (PBST) and heated in a microwave oven for 5 min to remove bound antibodies. Membranes were cooled, washed with PBST, reblocked with 5% nonfat milk, and reprobed overnight at 4 °C with rabbit monoclonal antibody to total AKT (Cell Signaling Technology, cat. no. 4691T; 1:1,000). Membranes were then incubated for 1 h at room temperature with HRP-conjugated goat anti-rabbit IgG (H+L) (Proteintech, cat. no. SA00001-2; 1:5,000). Signals were developed with enhanced chemiluminescence substrate (Thermo Fisher Scientific, cat. no. 32106) and imaged with a ChemiDoc MP system (Bio-Rad). AKT phosphorylation was quantified as the phospho-AKT-to-total AKT signal ratio; GAPDH served as an additional loading control.

### Flow cytometry of human B cells

Cells were washed in FACS buffer (PBS containing 2% FBS), stained with Zombie Aqua Fixable Viability Dye (BioLegend, cat. no. 423101), and incubated with antibodies for 30 min on ice in the dark. The 48-h activation panel comprised anti-CD69–FITC (clone FN50; BioLegend, cat. no. 310904), anti-CD80–APC (clone 2D10; BioLegend, cat. no. 305219), and anti-CD86–PE (clone BU63; BioLegend, cat. no. 374205). The day-7 memory-phenotype panel comprised anti-CD27–APC (clone LG.3A10; BioLegend, cat. no. 124212) and anti-human IgG Fc–PE/Cyanine7 (clone QA19A42; BioLegend, cat. no. 366908). Samples were acquired on a CytoFLEX cytometer (Beckman Coulter) after compensation with single-stained controls. Debris, doublets, and nonviable cells were excluded sequentially. Activation-marker expression was reported as mean fluorescence intensity among live singlets. CD27^+^IgG^+^ cells were reported as a memory-phenotype population rather than as functionally validated memory B cells. Data were analyzed in FlowJo v10. Representative gating strategies for CD27^+^IgG^+^ memory-phenotype cells and activation-marker analyses are shown in Fig. S8A and Fig. S8C, respectively.

### Quantification of secreted human IgG

Supernatants collected after 7 days of stimulation were clarified at 400g for 5 min and stored at −80 °C. Total human IgG was quantified using an ELISA kit (Abcam, cat. no. ab195215) according to the manufacturer’s protocol. Samples and kit standards were assayed in parallel, and concentrations were interpolated from a four-parameter logistic standard curve. Samples outside the validated range were reanalyzed after dilution.

### Production of GFP-reporter EBV

The production of recombinant GFP-EBV was adapted from our previous protocol(*61*). Specifically, GFP-EBV was produced from CNE2-EBV cells maintained in RPMI 1640 with 700 μg ml⁻¹ G418. When cultures reached approximately 80% confluence, lytic induction was stimulated with 20 ng ml⁻¹ 12-O-tetradecanoylphorbol-13-acetate (TPA; Beyotime, cat. no. S1819) and 5 mM sodium butyrate (Beyotime, cat. no. S1539). Following an 18-h induction, the medium was replaced with fresh complete RPMI 1640 containing 10% FBS. Viral supernatants were collected 72 h post-induction, clarified by centrifugation at 2,000 × g for 15 min at 4 °C, and filtered through a 0.45-μm polyethersulfone membrane. Virions were subsequently concentrated by ultracentrifugation at 50,000 × g for 3 h at 4 °C, resuspended in serum-free medium, aliquoted, and stored at −80 °C. The infectious titers were calculated as GRU/mL based on the frequency of GFP-expressing Raji cells following infection.

### GFP-based EBV neutralization assay

EBV neutralization assays were conducted as previously described(*61*). Briefly, heat-inactivated post-vaccination sera were initially diluted 1:100 and then serially diluted threefold and incubated with GFP-EBV for 2 h at 37 °C. Serum-virus mixtures were then inoculated onto either Akata cells to assess B-cell infection or HEK293 cells as an epithelial-cell model, with virus-only controls included on each plate. After a 1-h adsorption period, complete culture medium was added, and cells were cultured for 24–48 h. GFP positivity was measured by flow cytometry (CytoFLEX). Relative infection (%) was calculated as: (percentage of GFP+ cells in serum-treated wells) / (mean percentage of GFP+ cells in virus-only wells) × 100. Relative neutralization (%) was defined as 100 minus relative infection. The relative neutralization values shown in Fig. 4D and E correspond to the initial 1:100 serum dilution and, rather than IC50 titers, were used for between-group comparisons.

### Cryo-EM sample preparation and data acquisition

For cryo-EM sample preparation, 4 μL of the CB7t3-CR2 complex was applied to glow-discharged R2/1 Au 300 mesh Quantifoil grids. Grids were blotted for 4 s and flash-frozen in liquid ethane cooled by liquid nitrogen using a Vitrobot Mark IV (Thermo Fisher Scientific Inc.) operated at 4 °C and 100% humidity. The grids were subsequently transferred to a Titan Krios electron microscope (Thermo Fisher Scientific Inc.) operated at 300 kV and equipped with a Gatan K3 Summit direct electron detector and a GIF Quantum energy filter. Movie stacks were automatically collected using EPU with a preset defocus ranging from -1.2 μm to -2.0 μm in super-resolution mode. Data collection was performed at a nominal magnification of 130,000× for the CB7t3-CR2 complex, with a pixel size of 0.66 Å/pixel. The slit width on the energy filter was 20 eV, and the total dose was about 50 e^−^/Å^2^ for each micrograph stack.

### Image processing

Image processing was carried out in cryoSPARC(*62*) and RELION(*63*), and the strategies were shown in Fig. S9. The movie stacks were motion-corrected using MotionCor2(*64*), and the defocus values were estimated with Patch CTF estimation. Micrographs with contaminations or the maximum resolution lower than 6 Å were excluded from calculation, resulting in a total of 8,024 micrographs for structure determination.

A total of 346,812 particles were auto-picked using blob picker, extracted with a box size of 300 pixels from 500 micrographs, and classified into 50 classes by 2D classification. Representative 2D class averages were selected as the training dataset for Topaz Train(*65*). Then, 1,887,172 particles were auto-picked from 8,024 micrographs using Topaz. After one round of 2D classification, particles were selected and then subjected to Non-uniform Refinement with C3 symmetry(*66*). The particles were then imported to RELION and followed by 3D classifications with Relaxed C3 symmetry.

One density map with 384,769 particles was selected, which displays the best 3D features and resolution. Next, the selected particles were used to perform a 3D Refinement and yielded a reconstruction with a resolution of 3.1 Å. To better resolve the density, the particles were imported to cryoSPARC and subjected to Local Refinement, a final 2.97 Å cryo-EM map was obtained. All reported resolutions were estimated based on the gold-standard FSC 0.143 criterion.

### Model building and refinement

To generate an initial model, the AlphaFold3-predicted structure was docked into the cryo-EM density maps using Chimera(*67*). The models were iteratively rebuilt in COOT and refined in Phenix(*68*, *69*). The statistics for the cryo-EM data collection and model refinement were reported in supplemental Table 1. The refined coordinates and cryo-EM data were deposited into PDB and Electron Microscopy Data Bank, respectively.

### Mouse immunization and sample collection

Two parallel immunization cohorts were established, with *n* = 4 mice per group. In the non-adjuvanted cohort, mice received PBS or the indicated immunogen at a dose containing the same molar amount of gHgL as 2 μg of monomeric gHgL. In the alum-adjuvanted cohort, mice received Imject Alum Adjuvant alone or the corresponding equimolar immunogen dose formulated with Imject Alum Adjuvant (Thermo Fisher Scientific, cat. no. 77161). The mass of each immunogen was adjusted according to its molecular mass and the number of gHgL copies per assembly to provide an equal gHgL-molar dose across vaccine groups. Antigen and adjuvant were mixed at a 1:1 volume ratio by gentle end-over-end rotation for 30 min at room temperature immediately before administration. For the adjuvant-only control, an equivalent volume of PBS was mixed with adjuvant using the same procedure. Formulations were administered subcutaneously in a total volume of 100 μl per mouse at weeks 0, 3, and 6. Serum was collected at weeks 2, 5, and 8. Blood collected for serum preparation was allowed to clot for 1 h at room temperature, centrifuged at 12,000 rpm for 5 min, and stored at −80 °C. At week 8, mice were euthanized by carbon dioxide inhalation in accordance with institutional procedures. Peripheral blood for cellular analysis was collected into EDTA-containing tubes, one inguinal lymph node per mouse was collected for cryosection immunofluorescence, and spleens were collected for flow-cytometric analysis. Body weight and general clinical condition were monitored throughout the study.

### Serum anti-gHgL and anti-mC3d ELISA

High-binding 96-well microplates (Corning) were coated overnight at 4 °C with purified gHgL at 20 μg ml⁻¹ in PBS to measure anti-gHgL IgG. For anti-mC3d IgG assays, plates were coated under the same conditions with purified mouse C3d at 50 μg ml⁻¹. Plates were washed with PBS containing 0.05% Tween-20 (PBST) and blocked with 5% (w/v) BSA in PBST for 1 h at 37 °C. Mouse sera were diluted initially 1:100 in blocking buffer and then serially diluted threefold. Diluted sera were incubated on the plates for 1 h at 37 °C. After washing, bound IgG was detected with HRP-conjugated goat anti-mouse IgG (H+L) (Absin, cat. no. abs20001; 1:5,000) for 30 min at 37 °C. Plates were developed with TMB substrate (Tiangen Biotech) for 5–10 min, and reactions were stopped with 2 M H₂SO₄. Absorbance was measured at 450 nm with reference subtraction at 620 nm on an Epoch 2 microplate reader (BioTek). Binding-response area under the curve (AUC) was calculated from background-corrected absorbance values across the serum dilution series using the same dilution points for all samples within an assay. For anti-gHgL assays, AUC values were log₁₀-transformed before statistical analysis and are reported as Log AUC. For anti-mC3d assays, AUC was calculated with log₁₀ reciprocal serum dilution as the x-axis and was not further log-transformed; these values are reported as AUC. Binding curves and summary AUC values were analyzed separately for the non-adjuvanted and alum-adjuvanted cohorts.

### Collection and processing of mouse blood and spleens

Spleens were placed immediately in cold RPMI 1640 medium and mechanically dissociated through 70-μm cell strainers to generate single-cell suspensions. Splenic erythrocytes were lysed with sterile ACK lysis buffer (Solarbio, cat. no. R1013) for 5 min at room temperature. Peripheral-blood lymphocytes were isolated from EDTA-anticoagulated blood by density-gradient centrifugation using a Mouse Peripheral Blood Lymphocyte Isolation Solution Kit (Solarbio, cat. no. P8620) according to the manufacturer’s instructions. Cells were washed by centrifugation at 400g for 10 min, resuspended in FACS buffer, and counted with an automated cell counter. Approximately 1 × 10⁶ viable cells were used for each staining panel. Equal numbers of input cells were stained, and the same total number of cellular events was acquired for samples compared within an experiment. Accordingly, count values represent gated events under standardized acquisition conditions rather than absolute cell numbers per spleen or per unit blood volume.

### Flow cytometry of mouse lymphocytes

Separate multicolor panels were used to quantify GC B cells, Tfh cells, gHgL-binding IgG^+^ memory-phenotype B cells and additional memory B cell subsets, and memory T cell subsets. Splenocytes were used for GC B cell, Tfh cell, additional memory B cell subset, and CD4^+^/CD8^+^ memory T cell analyses. Peripheral-blood lymphocytes were used for the gHgL-binding IgG^+^ memory-phenotype B cell analysis shown in Fig. 4J, whereas splenocytes were used for the corresponding analysis shown in Fig. S17. Samples were stained with Zombie Aqua Fixable Viability Dye (BioLegend, cat. no. 423101) and the indicated fluorochrome-conjugated antibodies for 30 min on ice. Antibodies included B220/CD45R (clone RA3-6B2, cat. no. 103225), IgD (clone 11-26c.2a, cat. nos. 405730 or 405740), mouse IgG (Poly4053, cat. no. 405308), CD80 (clone 16-10A1, cat. no. 104726), CD73 (clone TY/11.8, cat. no. 127206), PD-L2 (clone TY25, cat. no. 107216), GL7 (clone GL7, cat. no. 144606), CD95/Fas (clone SA367H8, cat. no. 152608), CD3 (clone 17A2, cat. no. 100216), CD4 (clone GK1.5, cat. no. 100510), CD8α (clone 53-6.7, cat. no. 100744), CXCR5 (clone L138D7, cat. no. 145512), PD-1 (clone 29F.1A12, cat. no. 135210), CD44 (clone IM7, cat. no. 103040), and CD62L (clone MEL-14, cat. no. 104412); all antibodies were from BioLegend. BF750-gHgL was included as an antigen probe where indicated.

After sequential exclusion of debris, doublets, and dead cells, GC B cells were defined as B220^+^IgD^−^GL7^+^CD95^+^ cells. Tfh cells were operationally defined as CXCR5^+^ cells within live CD3^+^CD4^+^ T cells, and PD-1^+^ Tfh cells were defined as PD-1^+^ cells within the Tfh-cell gate. Class-switched antigen-binding memory-phenotype B cells were defined as B220^+^IgD^−^IgG^+^BF750-gHgL-binding cells. Additional memory B cell subsets were defined within B220^+^IgD^−^ cells by CD80^+^CD73^+^ or CD80^+^PD-L2^+^ coexpression. CD4^+^ and CD8^+^ central memory T cells were defined as CD44^+^CD62L^+^, and effector memory T cells were defined as CD44^+^CD62L^−^. Samples were acquired on a CytoFLEX cytometer and analyzed in FlowJo v10 using gates established with single-stained controls and appropriate biological negative controls. For panels displayed as gated event counts, gated events were compared after acquisition of the same total number of cellular events per sample. Representative gating strategies are shown for splenic GC B cells (Fig. S15A), splenic Tfh cells (Fig. S16A), and splenic gHgL-binding IgG^+^ memory-phenotype B cells (Fig. S17A); the latter marker hierarchy was also applied to the peripheral-blood analysis in Fig. 4J. Gating strategies for additional memory B-cell and memory T-cell subsets are shown in Fig. S21 and Fig. S22, respectively.

### Lymph node cryosection immunofluorescence

One inguinal lymph node from each mouse was embedded in optimal cutting temperature (OCT) compound, snap-frozen, and sectioned at 8 μm. Sections were air-dried for 30 min, fixed in ice-cold acetone for 10 min, and blocked with PBS containing 10% normal donkey serum for 1 h at room temperature. For the 4-h lymph-node trafficking experiment, sections were incubated overnight at 4 °C, protected from light, with Alexa Fluor 488–conjugated anti-mouse/human CD45R/B220 (clone RA3-6B2; BioLegend, cat. no. 103225; 1:50) and Alexa Fluor 700–conjugated anti-mouse CD21/CD35 (CR2/CR1) (clone 7E9; BioLegend, cat. no. 123432; 1:50). The mCherry signal from the injected fusion protein was detected directly without antibody amplification. For analysis of GC architecture at week 8, sections were stained overnight at 4 °C with Alexa Fluor 488–conjugated anti-B220 (clone RA3-6B2; BioLegend, cat. no. 103225; 1:50) and Alexa Fluor 647–conjugated anti-mouse/human GL7 antigen (clone GL7; BioLegend, cat. no. 144606; 1:50). Tfh localization was assessed in a single multiplex panel containing Alexa Fluor 488–conjugated anti-B220 (clone RA3-6B2; BioLegend, cat. no. 103225; 1:50), Alexa Fluor 700–conjugated anti-mouse CD4 (clone GK1.5; BioLegend, cat. no. 100430; 1:50), and PE/Dazzle 594–conjugated anti-mouse CD279/PD-1 (clone 29F.1A12; BioLegend, cat. no. 135228; 1:50). Sections were washed three times with PBS, counterstained with DAPI, and mounted with antifade mounting medium. Whole-section tiled images were acquired with identical laser power, detector gain, exposure, and acquisition settings for samples compared within an experiment. Images were processed in ImageJ/Fiji. For each complete lymph-node section, all GL7^+^ regions were segmented using a fixed threshold and summed to obtain the total GL7^+^ area per section (mm²). For B220/CD4/PD-1 images, CD4^+^PD-1^+^ double-positive regions were segmented using fixed channel thresholds and summed; the resulting area was divided by the total B220^+^ area in the same section and expressed as a percentage. When multiple nonadjacent sections were available from one mouse, section-level values for each endpoint were averaged to generate a single mouse-level value.

### *In vivo* draining lymph node targeting and *ex vivo* biodistribution

For acute trafficking studies, *n* = 4 mice per group received 10 μg of purified mCherry-fused CB7t3, mC3d, or 50A in 50 μl PBS by subcutaneous injection into the thigh; PBS-injected mice served as controls. Four hours later, mice were euthanized and the ipsilateral inguinal draining lymph nodes were excised. Lymph nodes were imaged *ex vivo* on an IVIS Spectrum system (PerkinElmer) using an mCherry-compatible filter set and an exposure selected to avoid signal saturation. Regions of interest were drawn around individual lymph nodes in Living Image 4.0, and average radiant efficiency was corrected for background. Cryosections from matched lymph nodes were used to assess follicular localization of the directly detected mCherry signal.

For whole-organ biodistribution, *n* = 4 mice received 10 μg of mCherry-fused CB7t3 by the same subcutaneous route. Four hours later, the heart, lungs, spleen, bilateral inguinal draining lymph nodes, kidneys, and liver were excised and imaged under identical acquisition settings. The two draining lymph nodes from each mouse were displayed as a bilateral pair. Organ fluorescence was interpreted relative to the draining lymph nodes and tissue background. The four mice shown in Fig. S20 were independent biological replicates.

### Statistical analysis and reproducibility

Statistical analyses were performed in GraphPad Prism 10. The biological replicate was an individual mouse or an independent human donor, as indicated in the figure legends. Technical replicates were averaged before biological-level analysis and were not treated as independent observations. For the histologic endpoints in Fig. S16D and E, section-level values were averaged within each mouse before group-level analysis, and group comparisons were performed using Kruskal–Wallis tests followed by Dunn’s multiple-comparisons tests. Data are presented as mean ± s.e.m. unless otherwise stated. Normality was assessed by the Shapiro–Wilk test and homogeneity of variances by Brown–Forsythe test. Multiple-group comparisons were performed by one-way ANOVA followed by Tukey’s multiple-comparisons test when assumptions were satisfied; otherwise, the Kruskal–Wallis test with Dunn’s multiple-comparisons test was used. Two-group comparisons used an unpaired two-tailed Student’s t test or Mann–Whitney U test as appropriate. P values are indicated using the significance thresholds specified in the figure legends. Nonparametric tests were used only where specified in the corresponding figure legend. Anti-gHgL AUC values were log₁₀-transformed before group comparison. Anti-mC3d AUC values in Fig. S19 were calculated over log₁₀ reciprocal serum dilution and analyzed without further transformation. Relative neutralization values in Fig. 4D and E were analyzed directly as percentages. All tests were two-sided, and P < 0.05 was considered statistically significant. Significance symbols, exact sample sizes, and the statistical test used for each panel are specified in the figure legends.

## Acknowledgement

This work was supported by grants from the Noncommunicable Chronic Diseases-National Science and Technology Major Project (2023ZD0501000), National Natural Science Foundation of China (U24A20743, 32441094, 82402614), Shenzhen Science and Technology Program (JCYJ20240813094918025), Young Talent Support Project of Guangzhou Association for Science and Technology (QT-2025-034), Fundamental Research Funds for Central Universities, Sun Yat-sen University (2026QNPY04), National Key Research and Development Program of China (2022YFC3400900), and Young Talents Program of Sun Yat-sen University Cancer Center (PT2227440003).

## Author contributions

Y.T.L., Y.N.Y., C.S. and M.S.Z. designed the work. Y.T.L. and G.L.B. performed protein design, purification and biochemical characterization. Y.N.Y., Y.T.L. and H.Z. performed B cell activation experiments, animal studies and related validation experiments. H.Q.X. and B.Z.C. performed cryo-EM data processing, structural determination and structural analysis. W.T.D. performed *in vivo* localization studies and sample collection from animal experiments. P.H.W. and C.X. performed virus neutralization assays. Y.T.L., Y.N.Y. and C.S. wrote the manuscript. C.S., Z.L. and M.S.Z. reviewed and revised the manuscript. All authors contributed to the completion of this manuscript and approved the manuscript submission.

## Data, code, and materials availability

Atomic coordinates of the designed CR2 binder–CR2 SCR1-2 complexes have been deposited in the Protein Data Bank (PDB) under accession code 43NF and in the Electron Microscopy Data Bank (EMDB) under accession code EMD-82007. All computational inputs, design protocols, scripts, and example notebooks used for CR2 binder generation would be publicly available. All computational designs and experimental materials generated in this study can be reproduced based on the methods described herein. Materials generated in this study are available from the corresponding authors upon reasonable request.

## Supplementary information

**Fig. S1.**
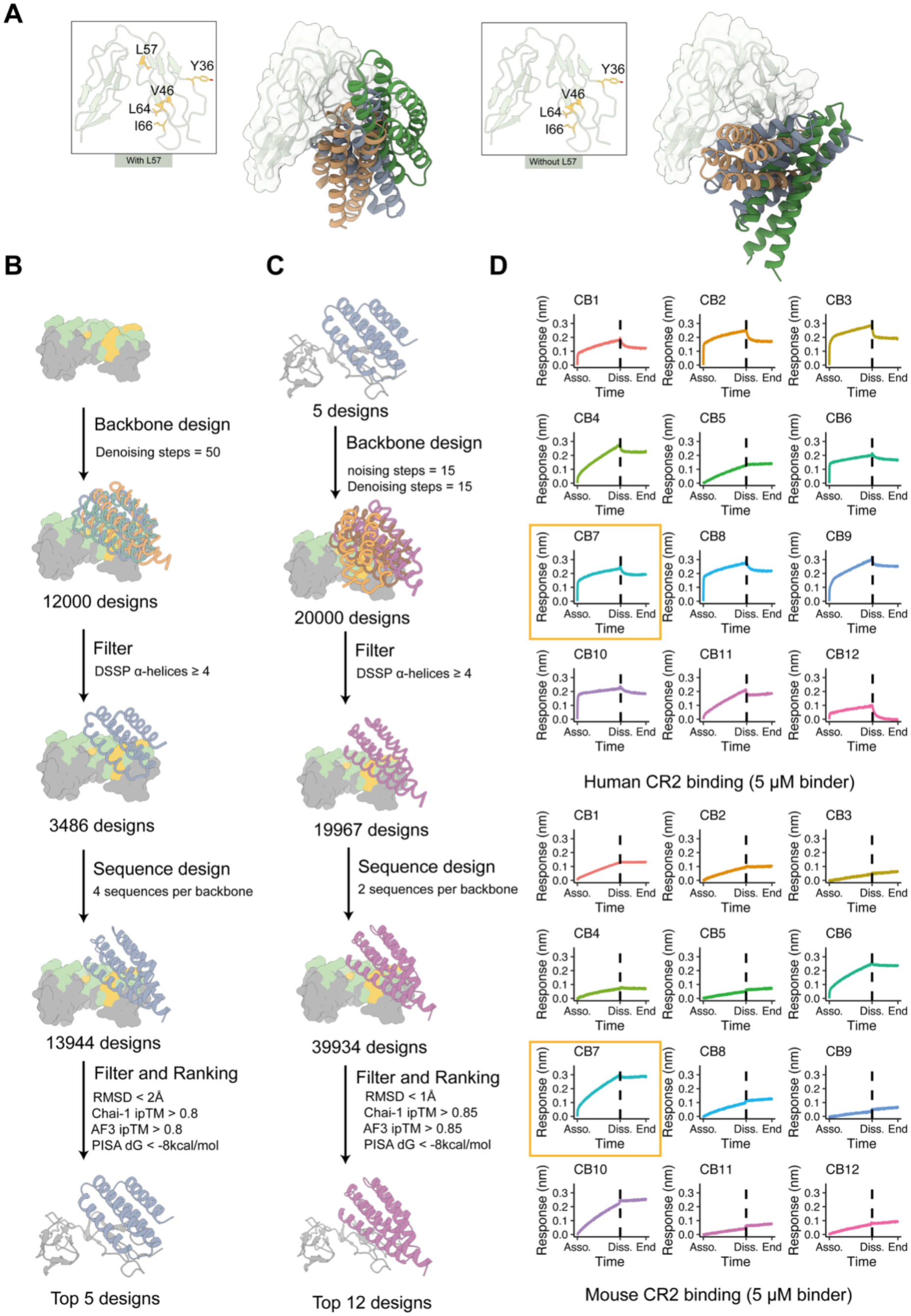
Design and optimization of monomeric CR2 binders. (A) Effect of including or excluding residue L57 as a hotspot constraint on the orientation of de novo-designed binders. (B) Workflow for de novo generation of monomeric CR2 binders. (C) Workflow for affinity optimization of CR2 binders. (D) Binding of 12 monomeric binders to human and mouse CR2 measured by BLI at a binder concentration of 5 μM.

**Fig. S2.**
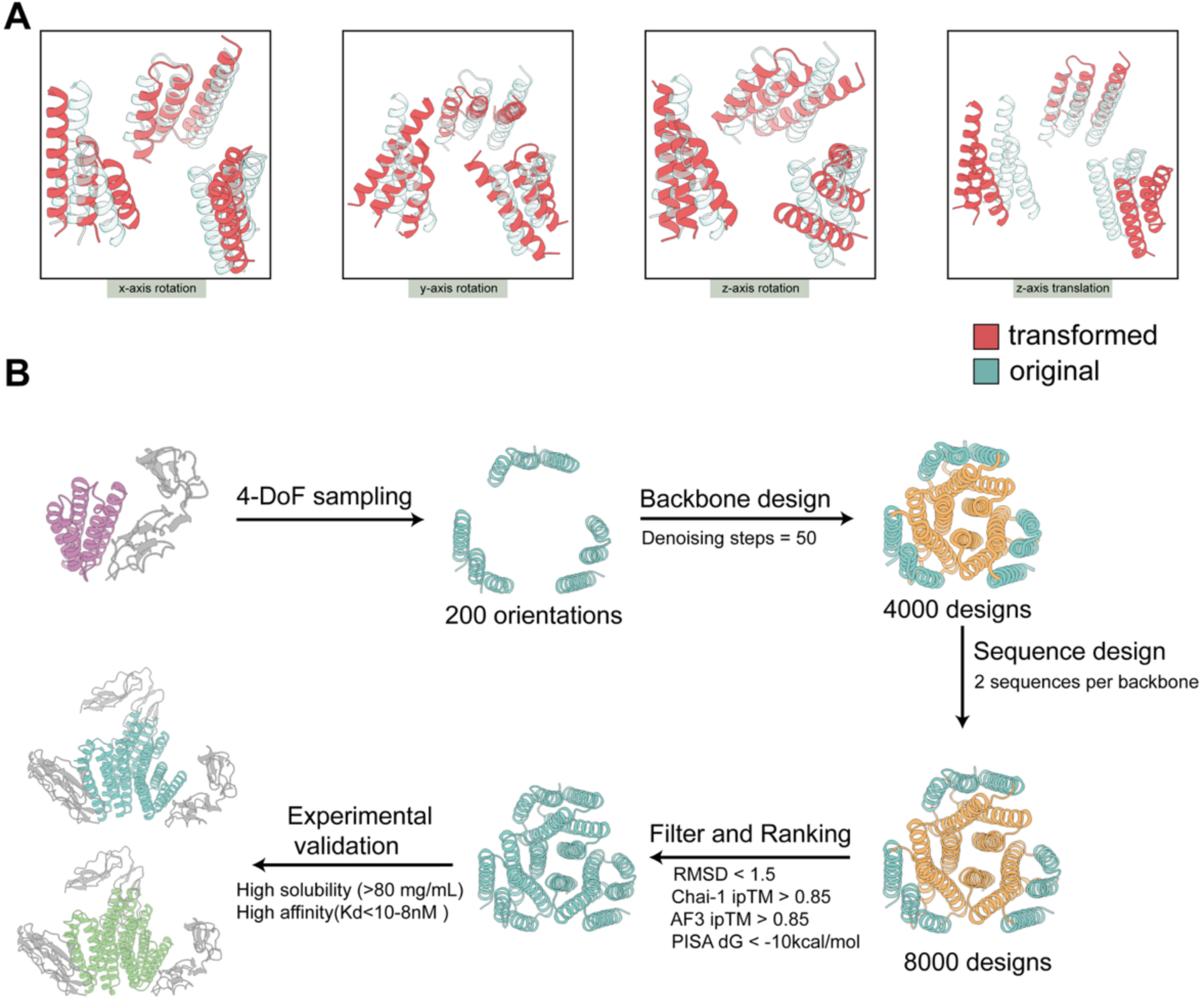
Design strategy for C3-symmetric trimeric binders. (A) Four degrees of freedom used for sampling C3-symmetric scaffold orientations. (B) Workflow for de novo design of C3-symmetric trimeric binder scaffolds.

**Fig. S3.**
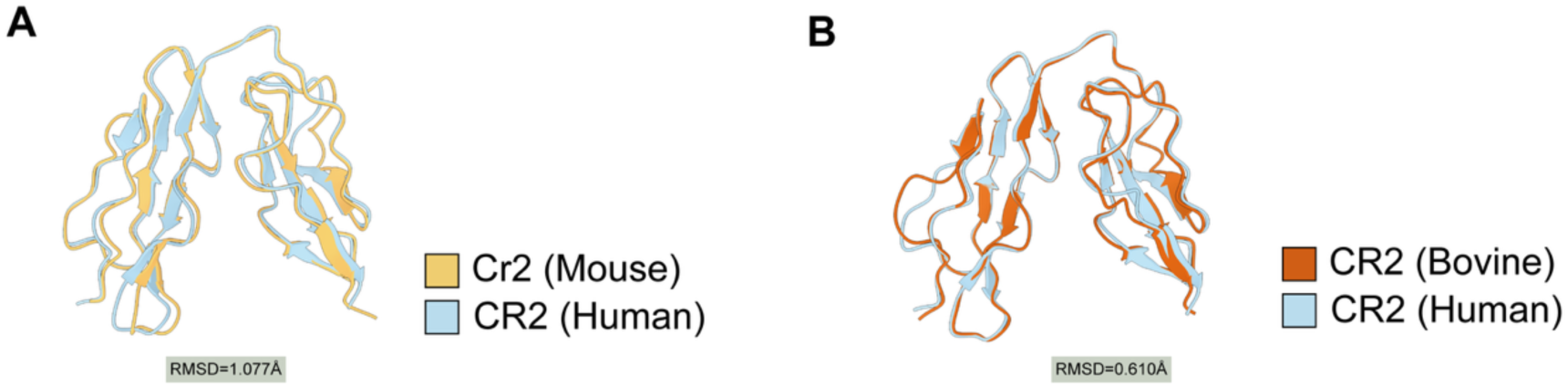
Structural comparison of CR2 orthologs predicted by AlphaFold3. (A) Structural superposition of human and mouse CR2 predicted by AlphaFold3 (RMSD = 1.077 Å). (B) Structural superposition of human and bovine CR2 predicted by AlphaFold3 (RMSD = 0.610 Å).

**Fig. S4.**
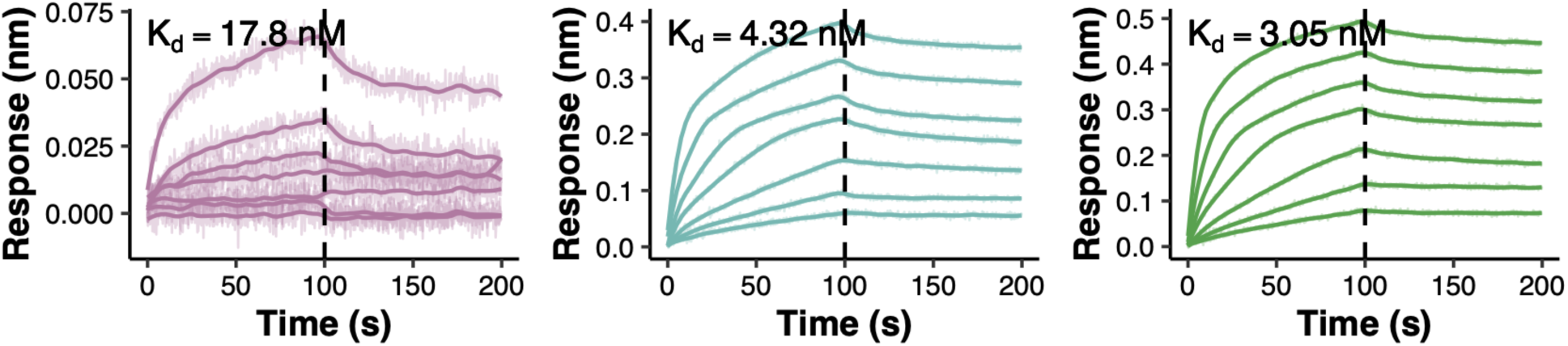
Binding kinetics of monomeric and trimeric CR2 binders toward mouse CR2. Binding kinetics of CB7, CB7t3, and CB7t4 toward mouse CR2 measured by BLI. The equilibrium dissociation constants (*K*_d_) are 17.8 nM for CB7, 4.32 nM for CB7t3, and 3.05 nM for CB7t4.

**Fig. S5.**
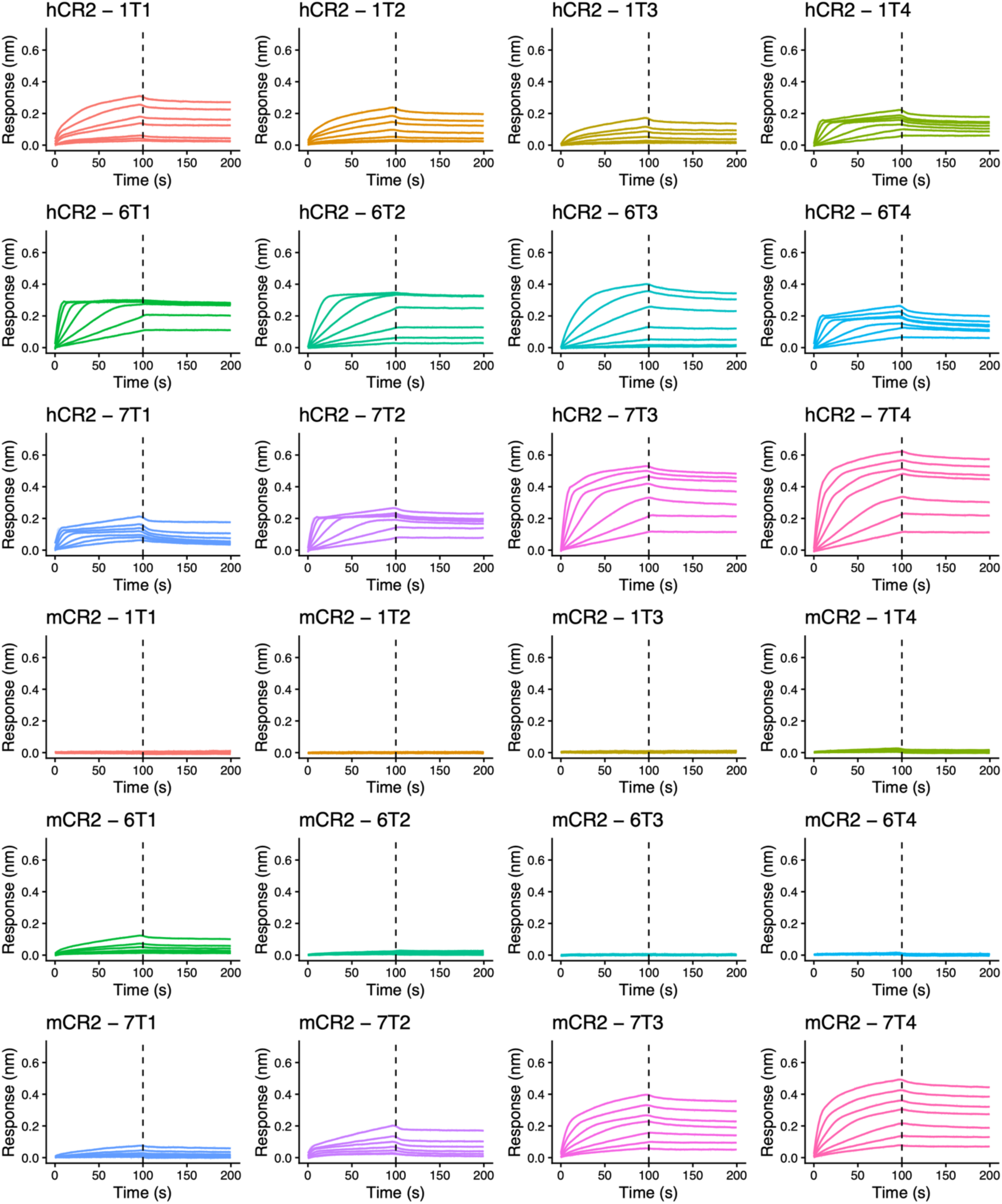
Binding characterization of trimeric CR2 binders. Binding kinetics of 12 trimeric binders toward human and mouse CR2 measured by BLI using a two-fold serial dilution series from a starting concentration of 400 nM (seven concentrations).

**Fig. S6.**
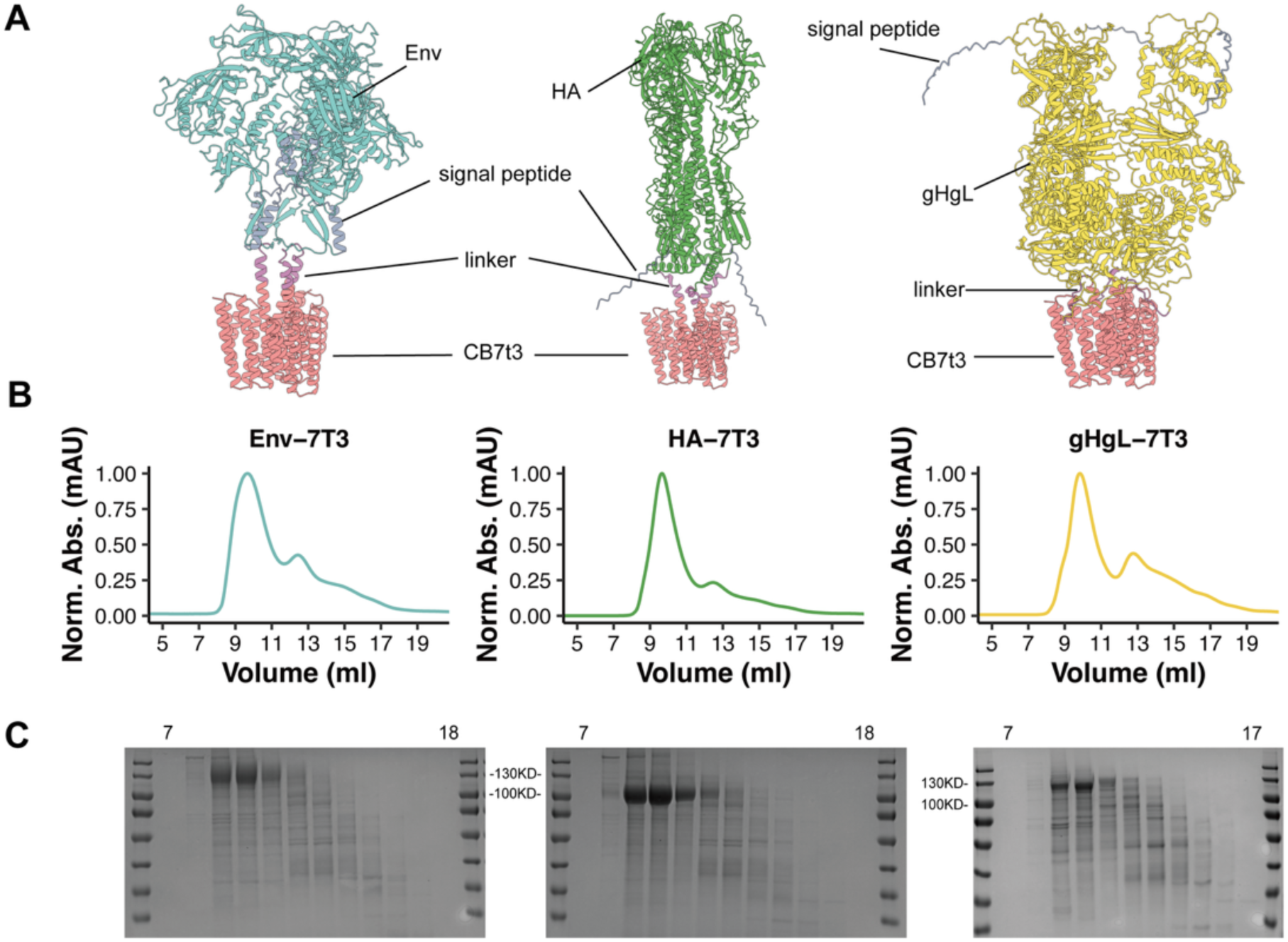
Fusion of CB7t3 to diverse antigens. (A) Structural models illustrating the spatial arrangement of different antigens fused to CB7t3. (B) SEC profiles of CB7t3 fusion proteins purified through the His-tag. (C) Coomassie-stained SDS–PAGE analysis of SEC fractions from purified CB7t3 fusion proteins.

**Fig. S7.**
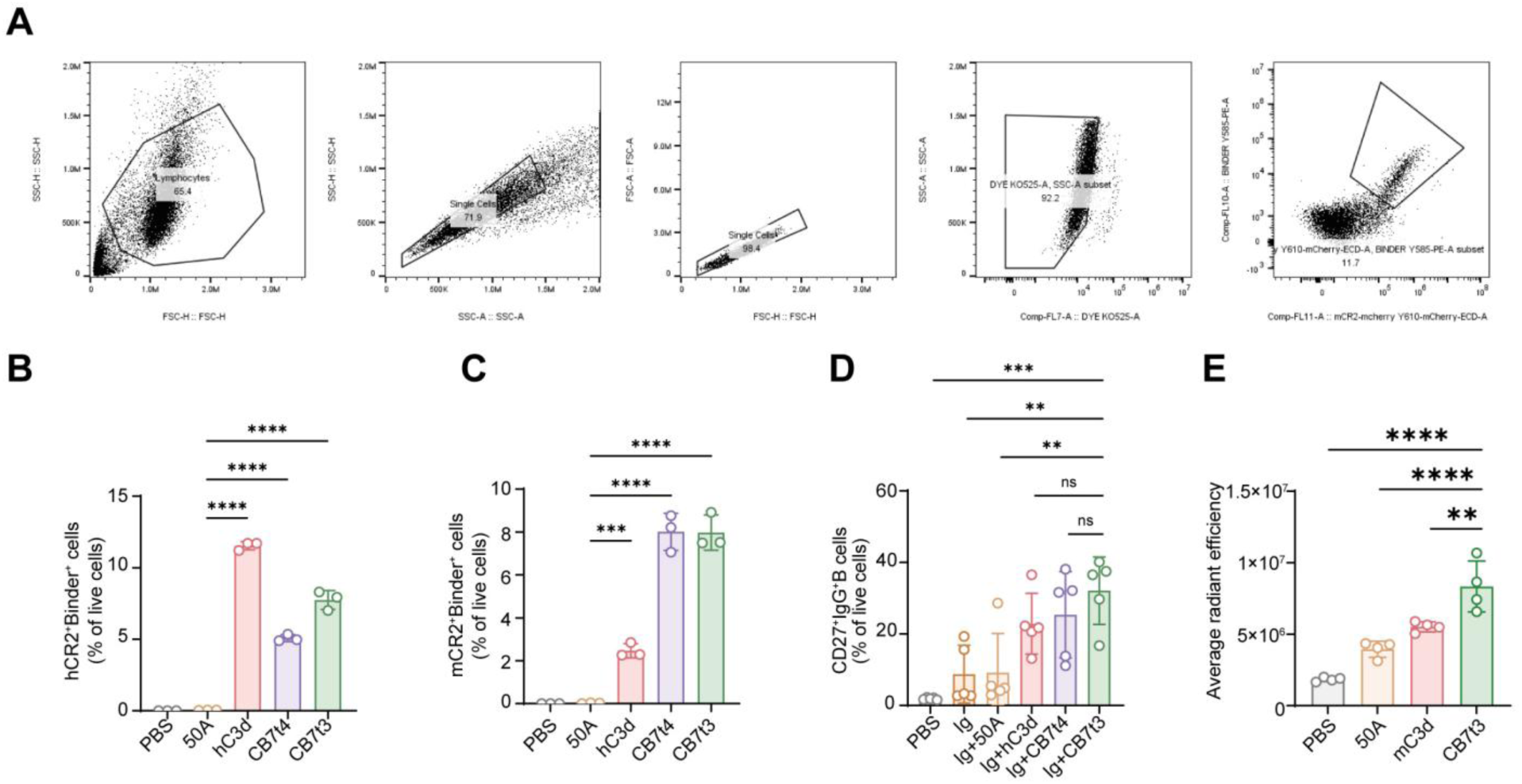
Designed CR2 trimers retain cross-species binding, enhance B-cell responses, and accumulate in draining lymph nodes. (A) Representative flow-cytometric gating strategy for identifying live mCherry-positive HEK293T cells after transient expression of human CR2 (hCR2) or mouse CR2 (mCR2) and for quantifying PE-labeled protein binding. (B and C) Percentages of live HEK293T cells double-positive for the mCherry CR2 reporter and PE-labeled binder after expression of hCR2 (B) or mCR2 (C) and incubation with PBS, PE-50A, PE-hC3d, PE-CB7t4, or PE-CB7t3. (D) Frequency of CD27^+^IgG^+^ memory-phenotype B cells among live primary human CD19^+^ B cells after 7 days of stimulation with low-dose anti-Ig alone or together with the indicated proteins. (E) Quantification of average radiant efficiency in excised draining inguinal lymph nodes 4 h after subcutaneous administration of mCherry-fused 50A, mC3d, or CB7t3; PBS-injected mice served as controls. Data in (B) and (C) are mean ± s.e.m. from three independent experiments (*n* = 3), data in (D) are mean ± s.e.m. from five independent experiments (*n* = 5), and data in (E) are mean ± s.e.m. from four mice per group (*n* = 4); each symbol represents one independent experiment in (B) to (D) or one mouse in (E). Statistical significance was determined by one-way ANOVA with Tukey’s multiple-comparisons test. \*\**P* < 0.01; \*\*\**P* < 0.001; \*\*\*\**P* < 0.0001; ns, not significant.

**Fig. S8.**
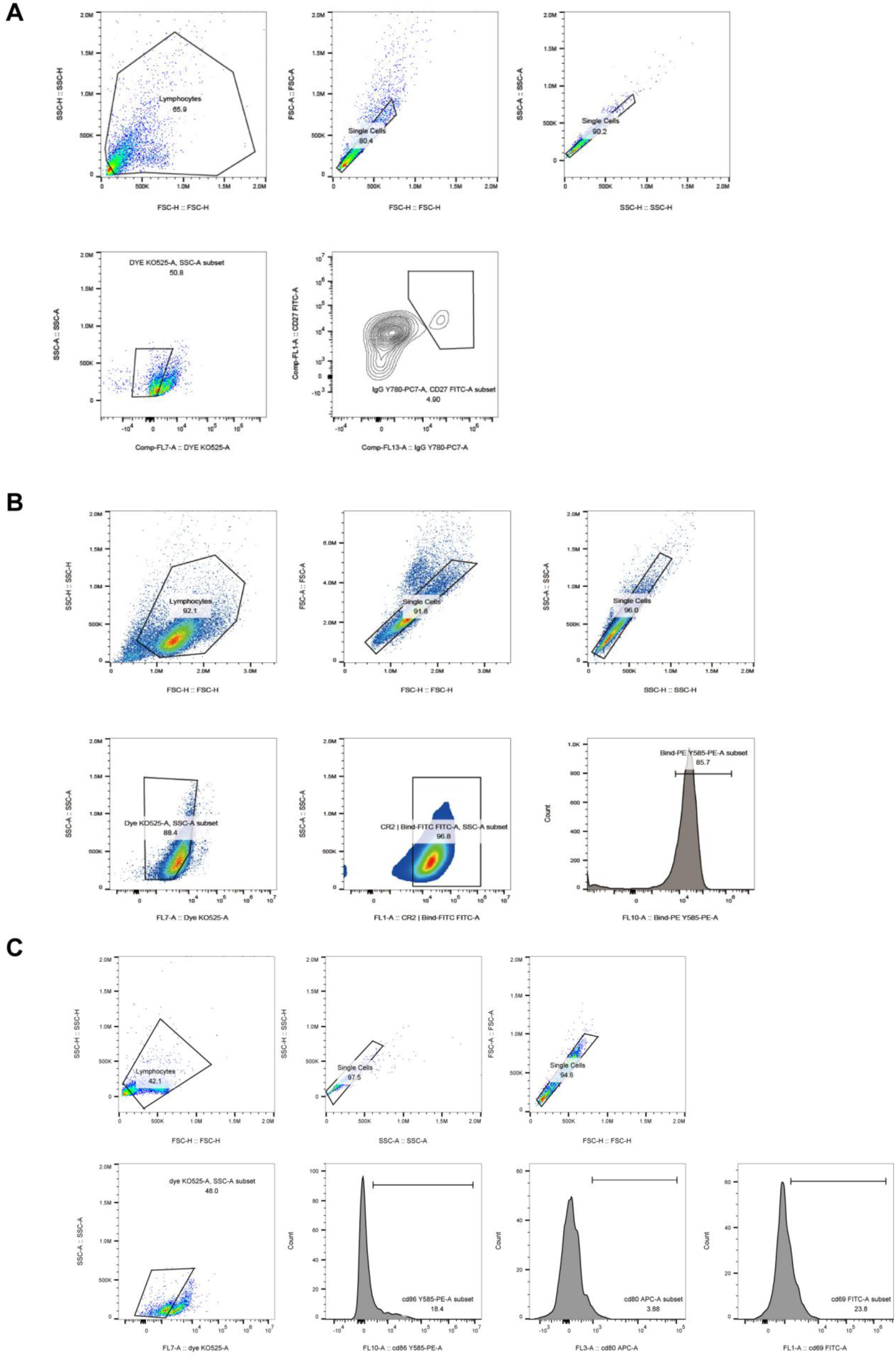
Flow cytometric gating strategies for the CR2-binding and B cell activation assays. (A) Gating strategy for CD27^+^IgG^+^ memory-phenotype B cells after sequential exclusion of debris, doublets, and dead cells. (B) Gating strategy for quantifying PE-labeled protein binding to endogenous CR2 on Akata and Raji B cell lines after sequential exclusion of debris, doublets, and dead cells. (C) Gating strategy for quantifying CD69, CD80, and CD86 expression among live singlet B cells. Representative plots are shown.

**Fig. S9.**
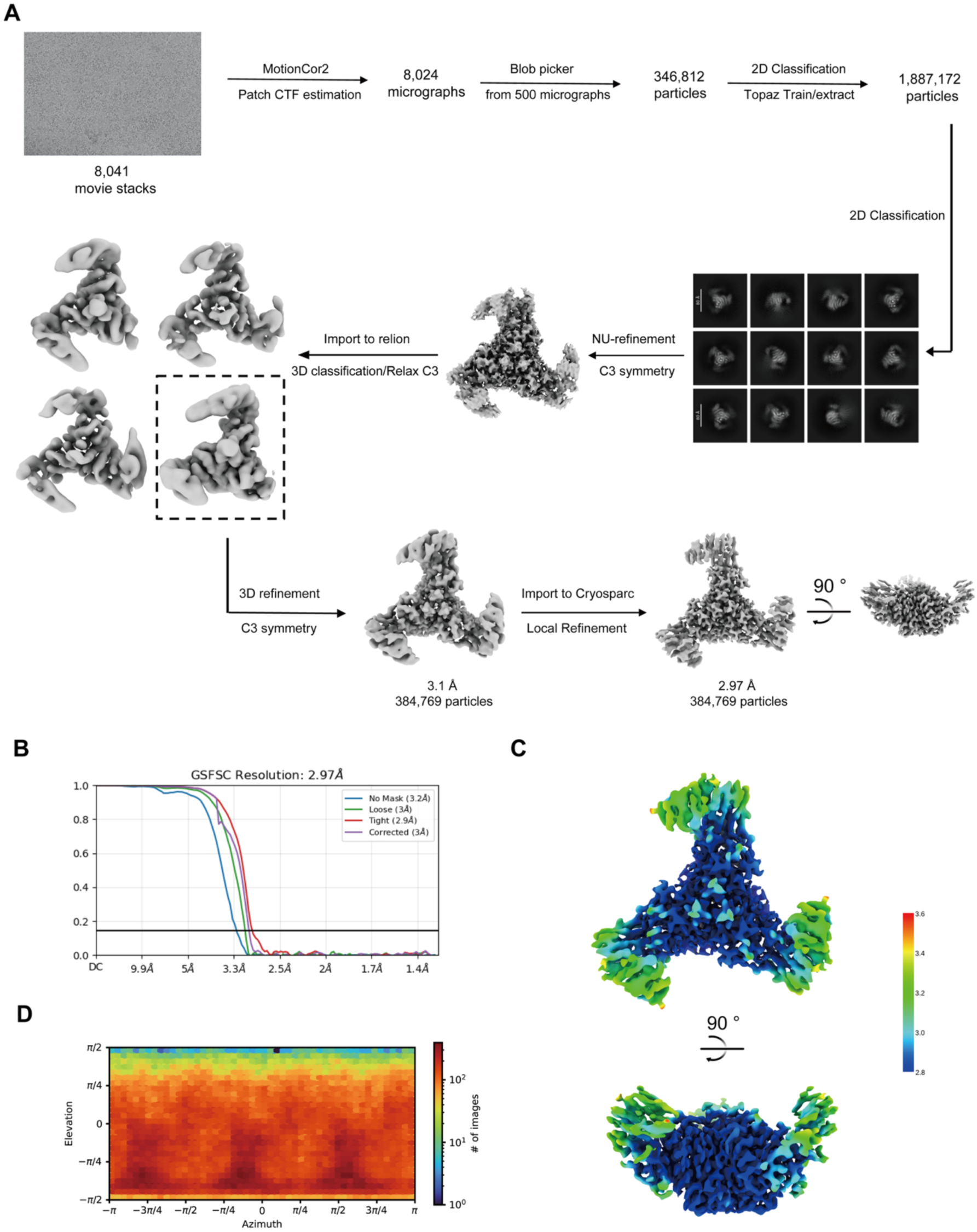
Cryo-EM data processing and validation of the CB7t3–CR2 complex. (A) Workflow for cryo-EM structure determination of the CB7t3–CR2 complex. (B) Gold-standard Fourier shell correlation (GS-FSC) curve indicating a final map resolution of 2.97 Å. (C) Local resolution estimation of the final density map. Colors indicate local resolutions ranging from 2.8 Å (blue) to 3.6 Å (red). (D) Particle orientation distribution used for three-dimensional reconstruction.

**Fig. S10.**
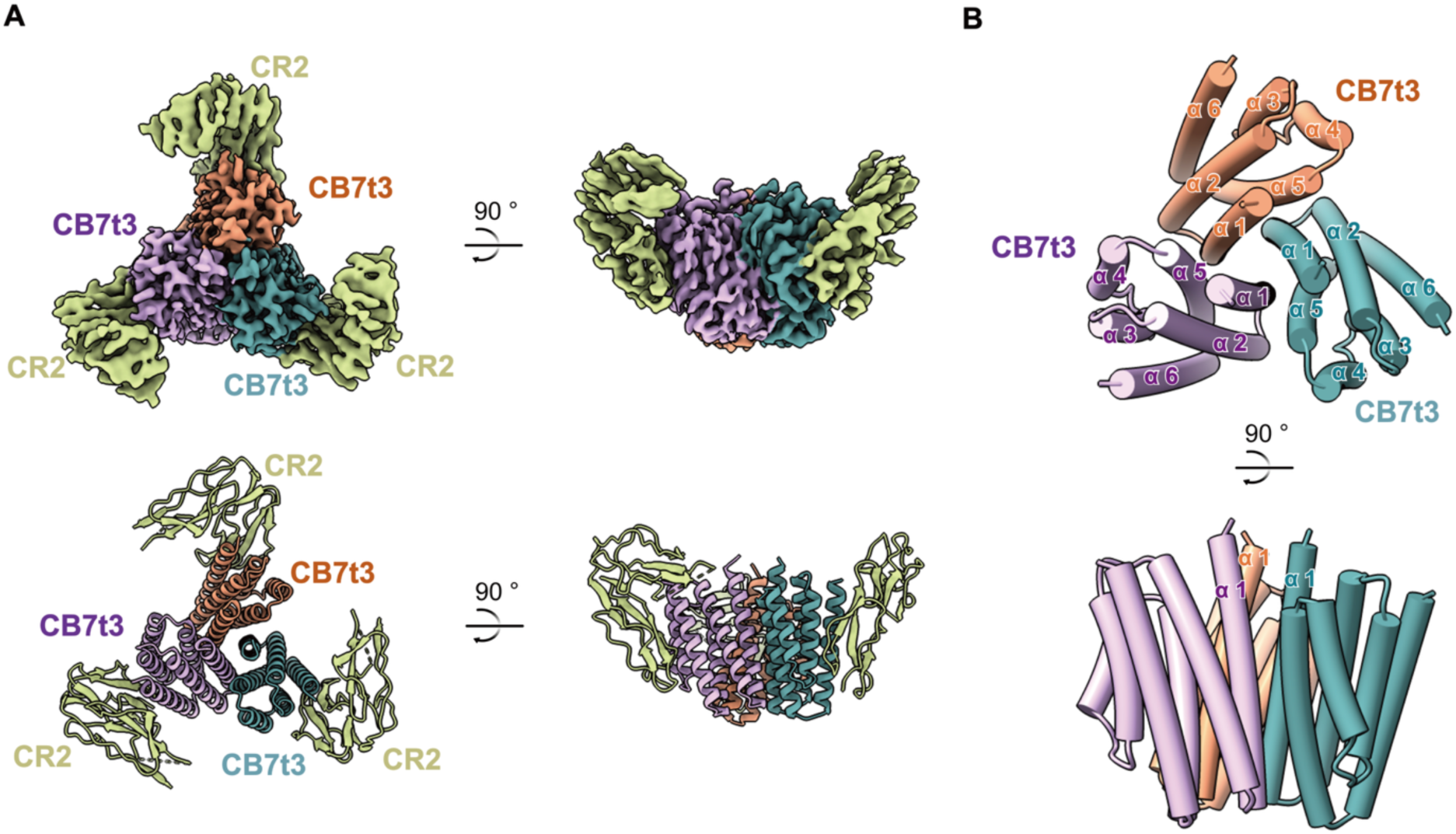
Overall architecture of the CB7t3–CR2 complex. (A) Top and side views of the cryo-EM density map together with the fitted atomic model. (B) Structural representation of the CB7t3 trimer, showing an architecture composed predominantly of α-helices.

**Fig. S11.**
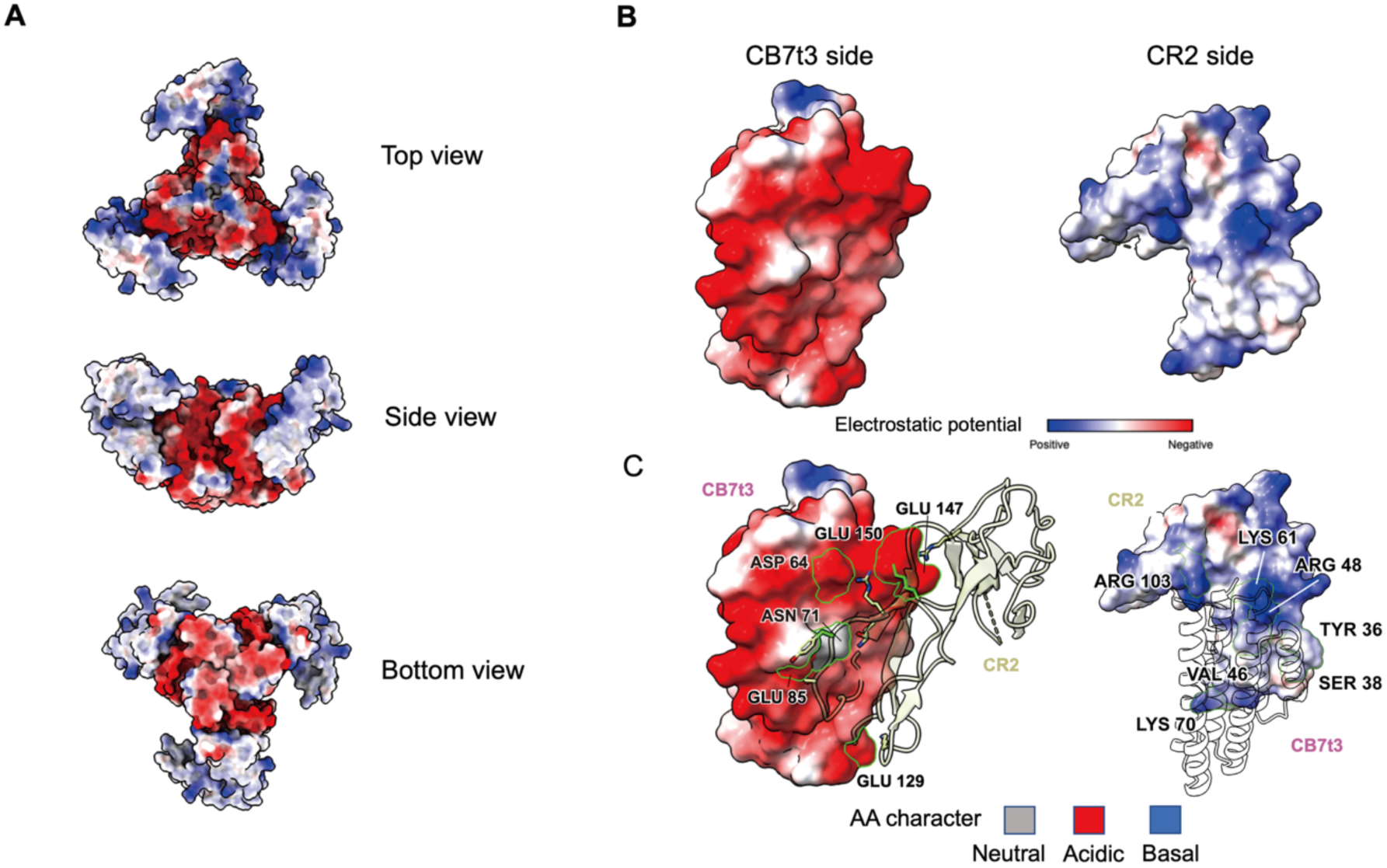
Electrostatic properties of the CB7t3–CR2 interface. (A) Electrostatic surface representation of the CB7t3–CR2 complex viewed from the top, side, and bottom. (B) Electrostatic complementarity at the CB7t3–CR2 binding interface. (C) Residues contributing to electrostatic interactions between CB7t3 and CR2.

**Fig. S12.**
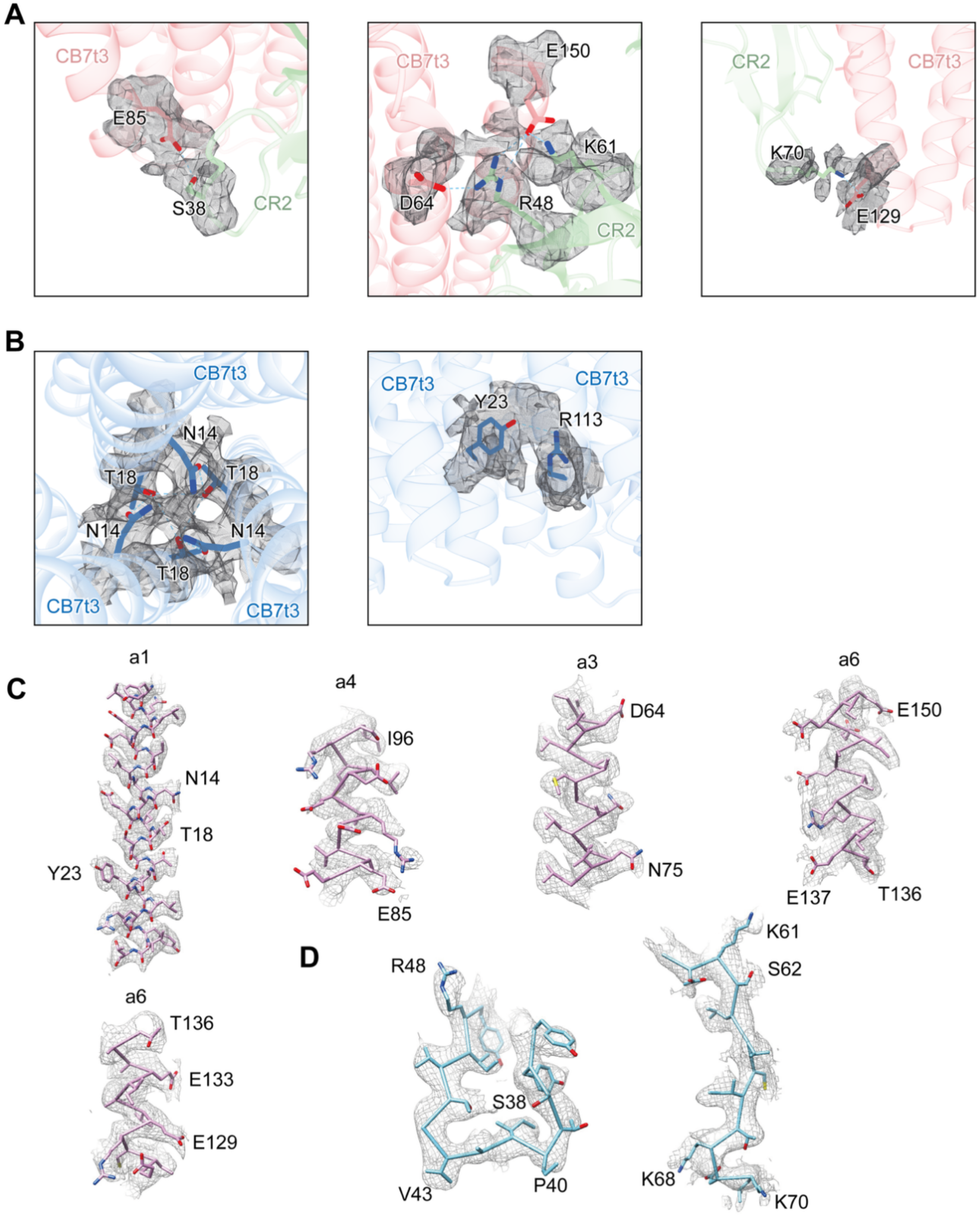
Cryo-EM density of key interface residues. (A) Cryo-EM densities corresponding to the side chains of key residues at the CB7t3–CR2 binding interface. (B) Cryo-EM densities corresponding to the side chains of key residues mediating intersubunit interactions within the CB7t3 trimer. (C) Local electron density maps around the CB7t3 hotspot residues (D) Local electron density maps around the CR2 hotspot residues

**Fig. S13.**
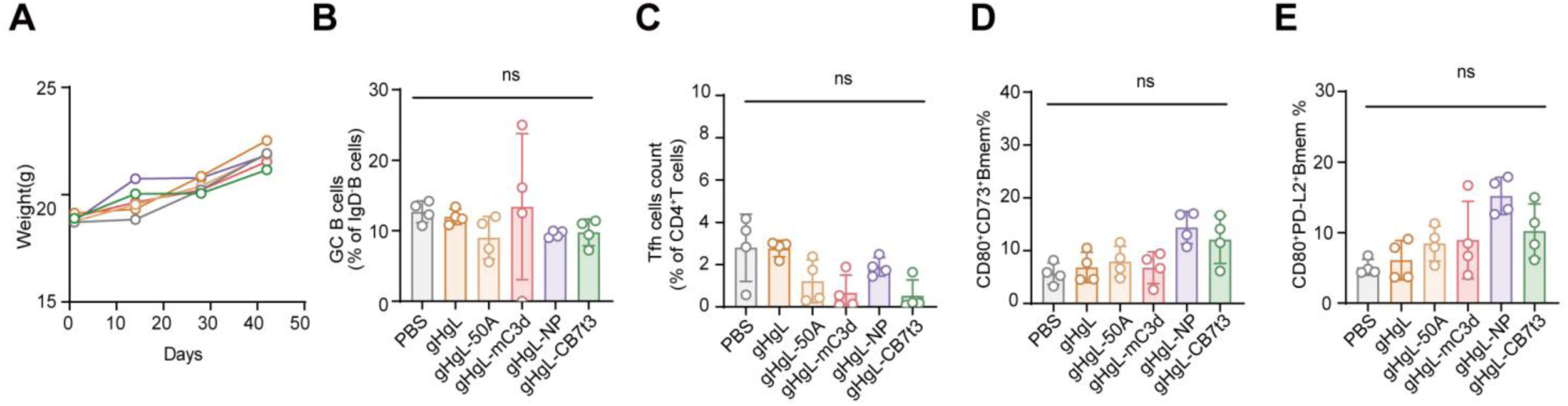
Cellular immune responses elicited by non-adjuvanted gHgL vaccine formulations. BALB/c mice (*n* = 4 per group) were immunized subcutaneously at weeks 0, 3, and 6 with the indicated non-adjuvanted formulations and analyzed at week 8, as in Fig. 4A. (A) Body weight trajectories during the immunization period. (B and C) Frequencies of splenic B220^+^IgD^−^GL7^+^CD95^+^ GC B cells among B220^+^IgD^−^ B cells (B) and splenic PD-1^+^ Tfh cells among CD3^+^CD4^+^ T cells (C) at week 8. (D and E) Frequencies of CD80^+^CD73^+^ (D) and CD80^+^PD-L2^+^ (E) memory B cell subsets among B220^+^IgD^−^ B cells. Data are mean ± s.e.m.; each symbol represents one mouse. Statistical significance was determined by one-way ANOVA with Tukey’s multiple-comparisons test; ns, not significant.

**Fig. S14.**
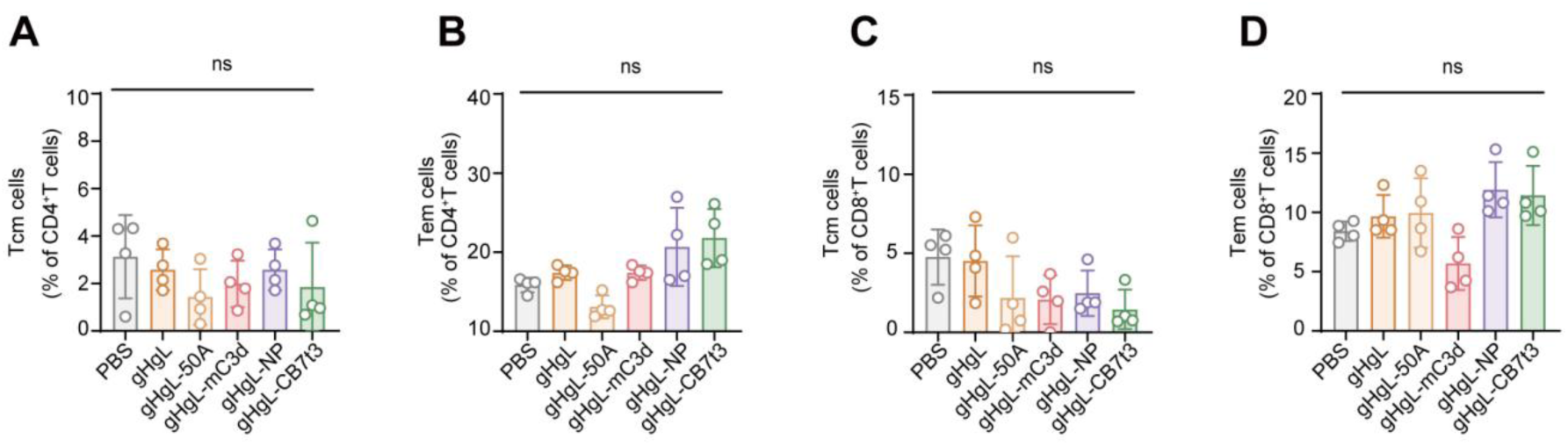
Memory T cell subsets after non-adjuvanted immunization. Mice (*n* = 4 per group) were immunized with the non-adjuvanted formulations shown in Fig. 4A, and splenic memory T cell subsets were analyzed at week 8. (A and B) Frequencies of CD4^+^ central memory T (Tcm) cells (CD44^+^CD62L^+^) (A) and CD4^+^ effector memory T (Tem) cells (CD44^+^CD62L^−^) (B). (C and D) Frequencies of CD8^+^ Tcm cells (C) and CD8^+^ Tem cells (D). Data are mean ± s.e.m.; each symbol represents one mouse. Statistical significance was determined by one-way ANOVA with Tukey’s multiple-comparisons test; ns, not significant.

**Fig. S15.**
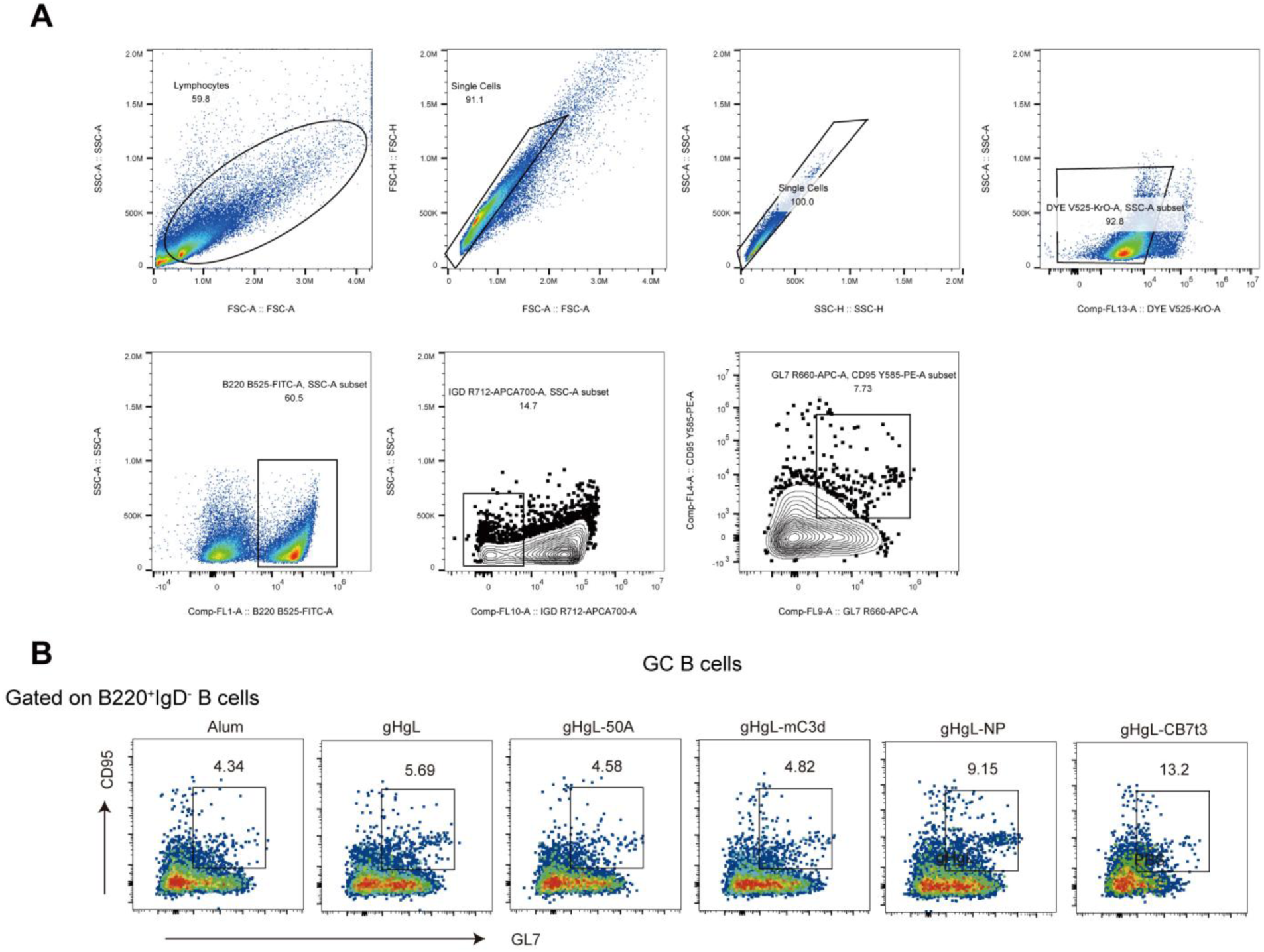
Flow-cytometric gating and representative plots for splenic GC B cells after alum-adjuvanted immunization. (A) Sequential gating strategy for splenic GC B cells. Lymphocytes and singlets were selected, dead cells were excluded, B220^+^ B cells were identified, and B220^+^IgD^−^ B cells were analyzed for GL7 and CD95 coexpression. (B) Representative GL7 versus CD95 plots gated on B220^+^IgD^−^ B cells from mice receiving alum alone or the indicated alum-adjuvanted gHgL formulations. Numbers indicate GC B cell frequencies within the parent population. Plots are representative of four mice per group.

**Fig. S16.**
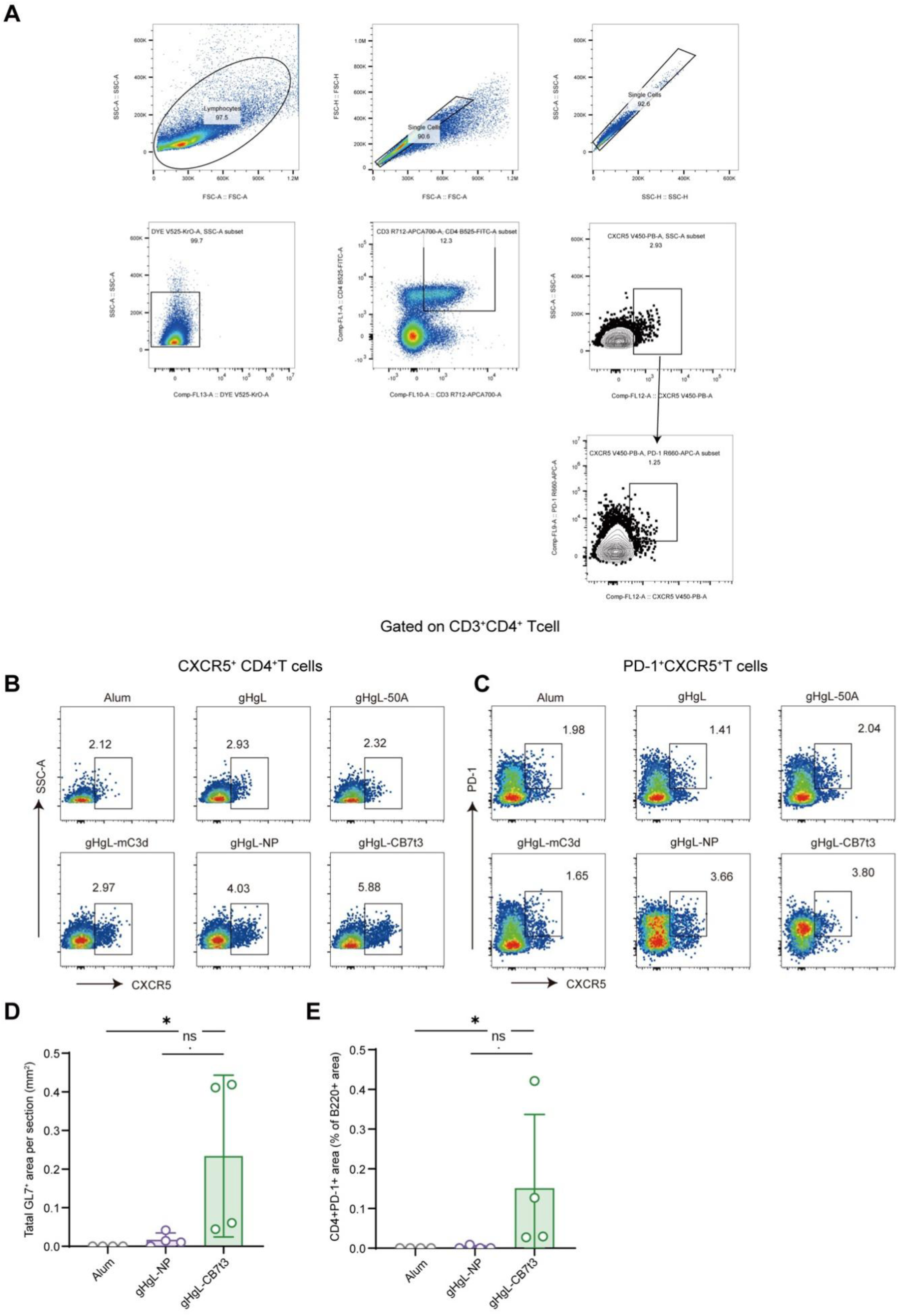
Flow-cytometric analysis of splenic Tfh cells and histologic quantification of draining lymph-node GL7^+^ area and CD4^+^PD-1^+^ area fraction after alum-adjuvanted immunization. (A) Sequential gating strategy for splenic Tfh cells. Lymphocytes and singlets were selected, dead cells were excluded, and CD3^+^CD4^+^ T cells were analyzed sequentially for CXCR5 and PD-1 expression. (B) Representative plots showing Tfh cells among CD3^+^CD4^+^ T cells in each immunization group. (C) Representative plots showing PD-1^+^ Tfh cells in each immunization group. Numbers indicate frequencies within the indicated parent populations. Plots are representative of four mice per group. (D) Total GL7^+^ area per complete draining lymph-node section at week 8 in selected groups representing the adjuvant control and the two highest-performing vaccine formulations: alum alone, gHgL-NP, and gHgL-CB7t3. (E) CD4^+^PD-1^+^ double-positive area expressed as a percentage of total B220^+^ area in matched complete draining lymph-node sections at week 8. Data are presented as mean ± s.e.m.; *n* = 4 mice per group, and each symbol represents one mouse. Group comparisons for (D) and (E) were performed using Kruskal–Wallis tests followed by Dunn’s multiple-comparisons tests. Adjusted P values are shown for the indicated comparisons; \**P* < 0.05; ns, not significant.

**Fig. S17.**
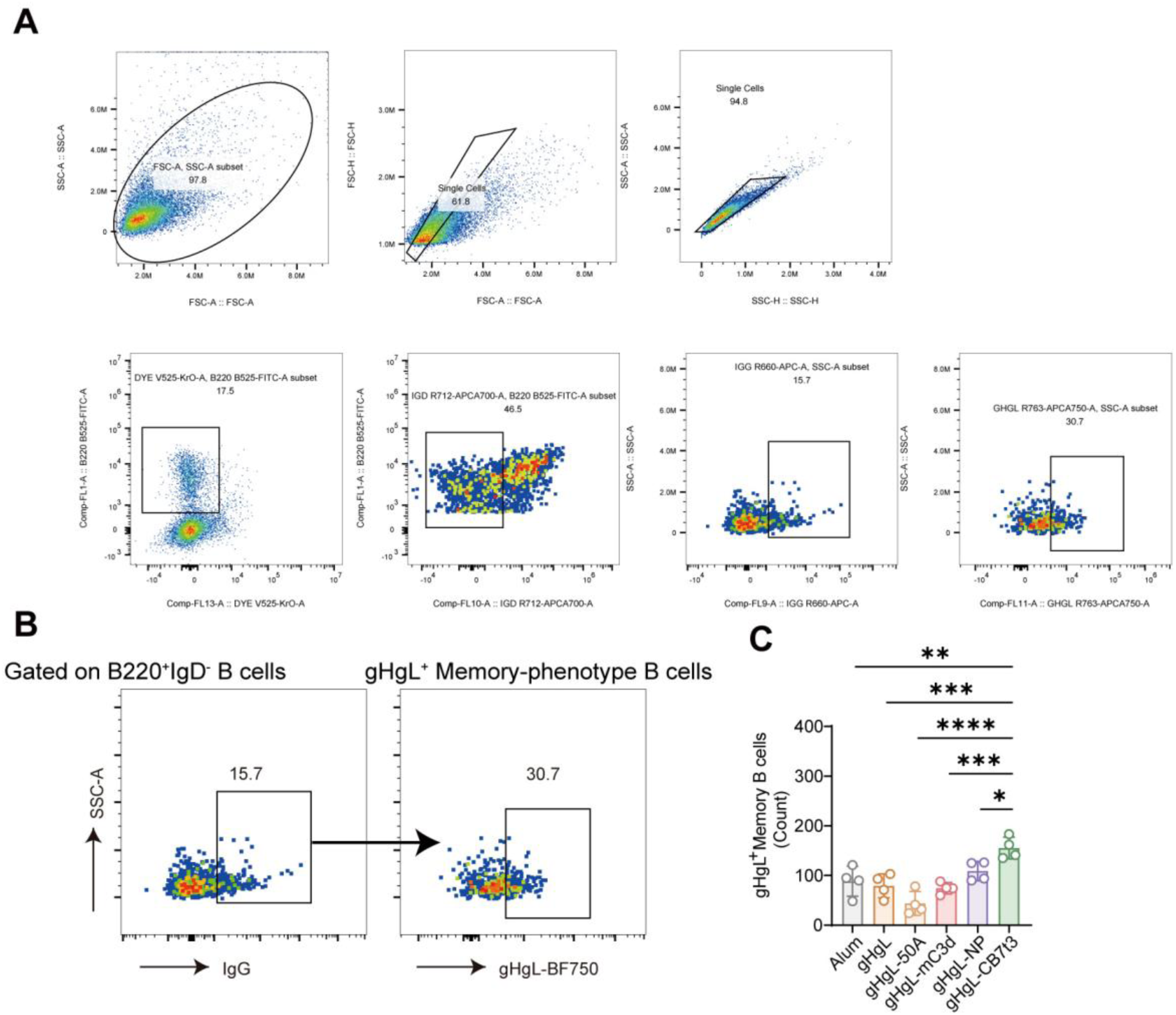
Flow-cytometric identification of splenic gHgL-binding IgG^+^ memory-phenotype B cells after alum-adjuvanted immunization. (A) Sequential gating strategy for splenocytes. After exclusion of debris, doublets, and dead cells, B220^+^IgD^−^ B cells were selected and analyzed for IgG expression and BF750-gHgL binding. (B) Representative plots showing the IgG^+^ gate among B220^+^IgD^−^ B cells and the subsequent BF750-gHgL-binding memory-phenotype B cell gate. (C) Cell count of gHgL-binding IgG^+^ memory-phenotype B cells in week-8 spleen samples from the indicated immunization groups. Equal numbers of input splenocytes were stained, and equal total cellular events were acquired; counts indicate cells within the terminal gate under standardized acquisition conditions, not absolute cell numbers per spleen. Data are presented as mean ± s.e.m.; *n* = 4 mice per group, and each symbol represents one mouse. Statistical significance was determined by one-way ANOVA with Tukey’s multiple-comparisons test. \**P* < 0.05; \*\**P* < 0.01; \*\*\**P* < 0.001; \*\*\*\**P* < 0.0001.

**Fig. S18.**
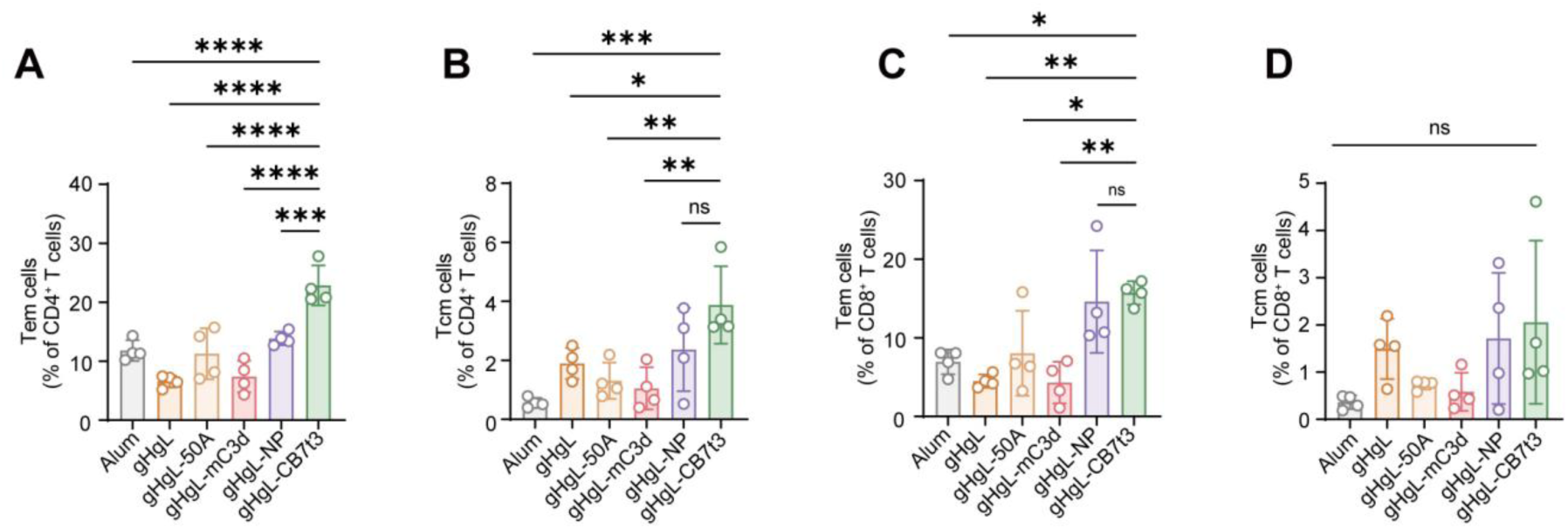
Splenic memory T cell responses after alum-adjuvanted immunization. Mice (*n* = 4 per group) were immunized with alum alone or the indicated alum-adjuvanted gHgL formulations, and splenic memory T cell subsets were analyzed at week 8. (A and B) Frequencies of CD4^+^ Tem cells (CD44^+^CD62L^−^) (A) and CD4^+^ Tcm cells (CD44^+^CD62L^+^) (B). (C and D) Frequencies of CD8^+^ Tem cells (C) and CD8^+^ Tcm cells (D). Data are mean ± s.e.m.; each symbol represents one mouse. Statistical significance was determined by one-way ANOVA with Tukey’s multiple-comparisons test. \**P* < 0.05; \*\**P* < 0.01; \*\*\**P* < 0.001; \*\*\*\**P* < 0.0001; ns, not significant.

**Fig. S19.**
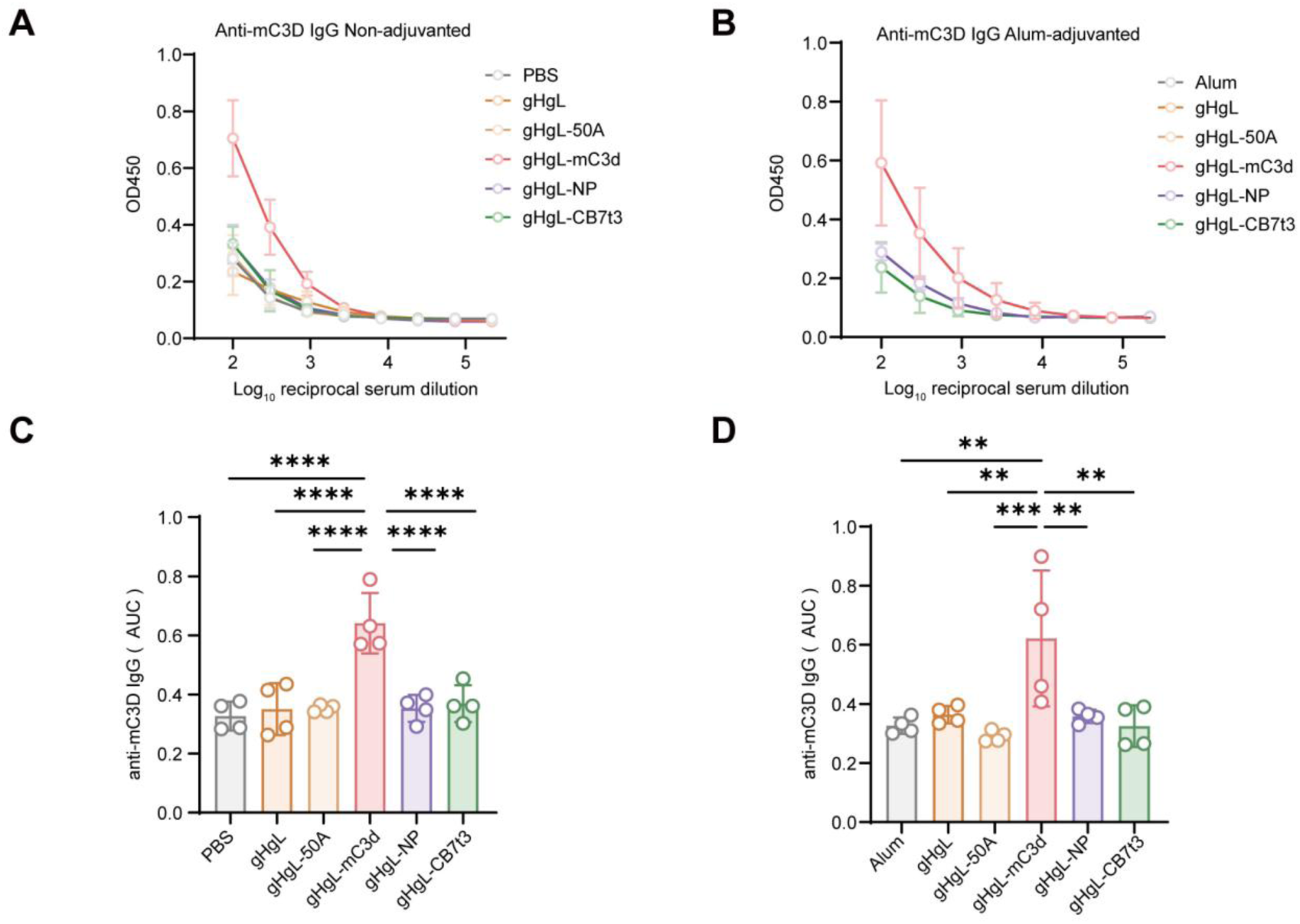
Serum anti-mC3d IgG responses after gHgL immunization. (A and B) Anti-mC3d IgG ELISA binding curves for week-8 sera from mice immunized with non-adjuvanted formulations (A) or alum-adjuvanted formulations (B). (C and D) Corresponding anti-mC3d IgG AUC values for the non-adjuvanted (C) and alum-adjuvanted (D) cohorts. AUC was calculated with log10 reciprocal serum dilution as the x-axis and was not further log-transformed. Data are mean ± s.e.m.; *n* = 4 mice per group, and each symbol represents one mouse. AUC values were compared by one-way ANOVA with Tukey’s multiple-comparisons test. \*\**P* < 0.01; \*\*\**P* < 0.001; \*\*\*\**P* < 0.0001.

**Fig. S20.**
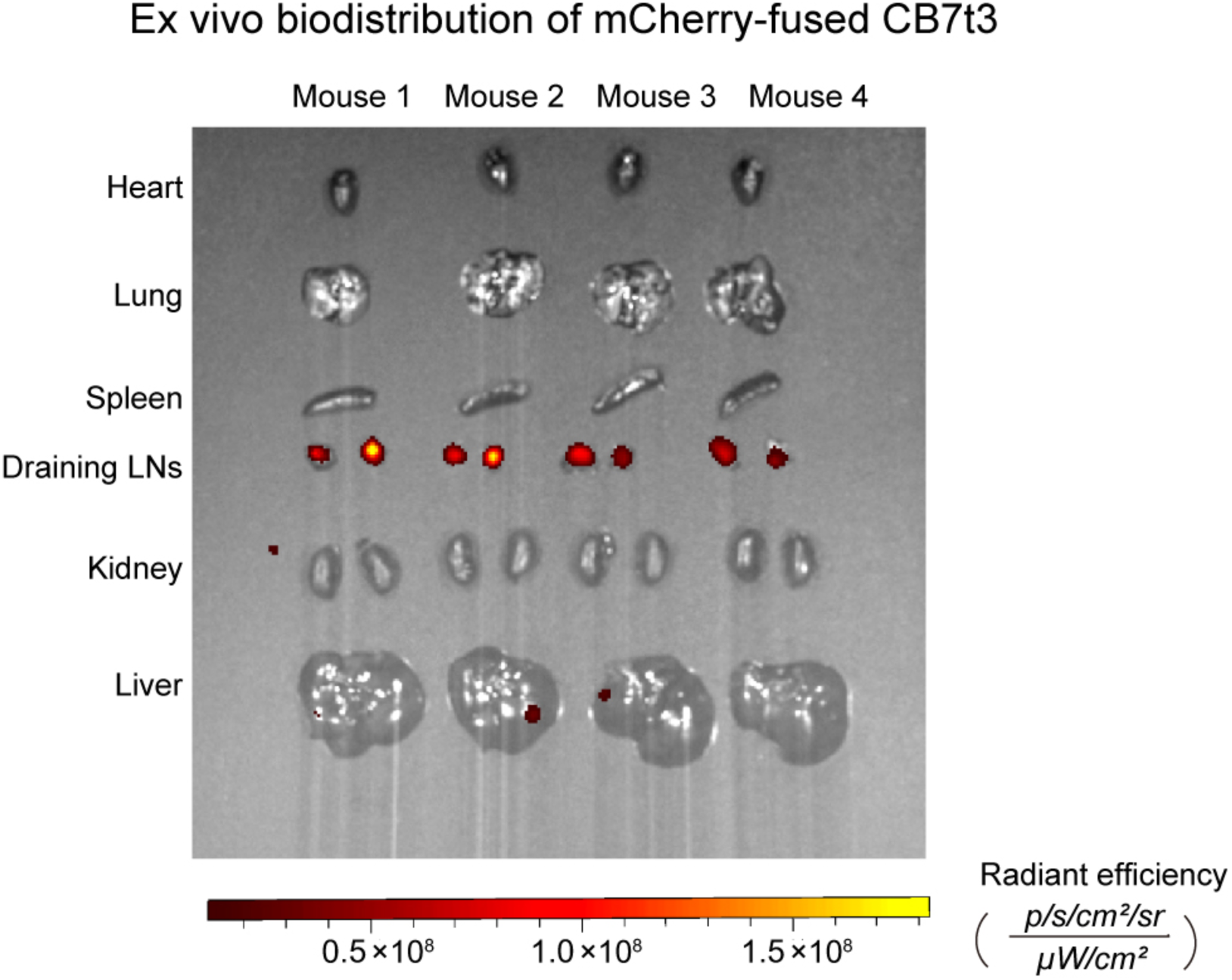
*Ex vivo* biodistribution of mCherry-fused CB7t3 after subcutaneous administration. *Ex vivo* mCherry fluorescence images of the heart, lungs, spleen, bilateral draining inguinal lymph nodes, kidneys, and liver collected from four mice 4 h after subcutaneous injection of mCherry-fused CB7t3. The two draining lymph nodes from each mouse are shown as a bilateral pair. Organs were imaged under identical acquisition settings, and the color scale indicates radiant efficiency.

**Fig. S21.**
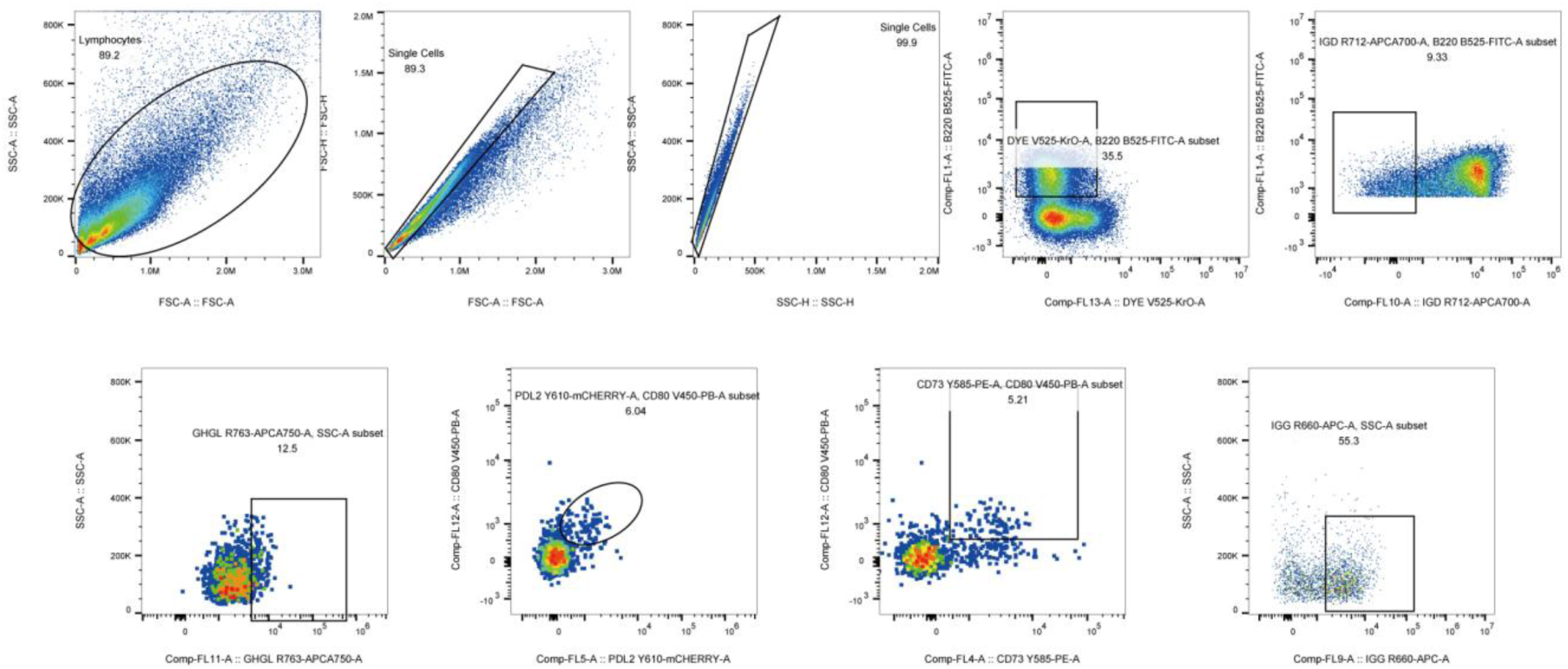
Flow-cytometric gating strategy for memory B cell phenotyping. Lymphocytes and singlets were selected, dead cells were excluded, and B220^+^IgD^−^ B cells were identified. Antigen-specific and class-switched B cells were quantified by BF750-gHgL binding and IgG expression, respectively; memory B cell subsets were defined by coexpression of CD80 with PD-L2 or CD73. Representative plots are shown from *n* = 4 mice per group.

**Fig. S22.**
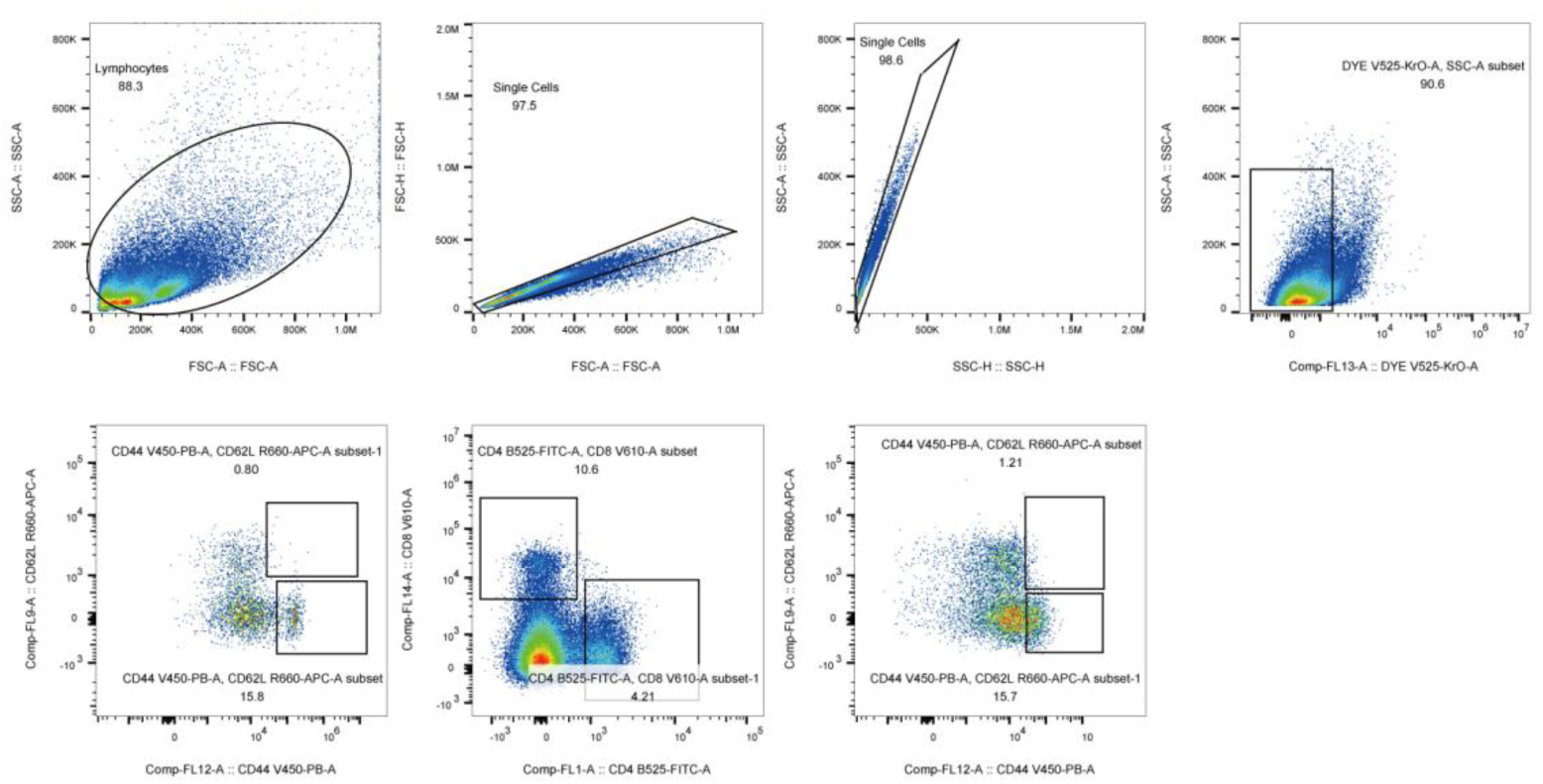
Flow-cytometric gating strategy for splenic CD4^+^ and CD8^+^ memory T cell subsets. After selection of lymphocytes, singlets, and live cells, CD4^+^ and CD8^+^ T cell populations were identified and analyzed for CD44 and CD62L expression. Tcm cells were defined as CD44^+^CD62L^+^, and Tem cells were defined as CD44^+^CD62L^−^. Plots are representative of four mice per group.

**Table S1.** Cryo-EM data collection, refinement and validation statistics.

|  |  |
| --- | --- |
|  | CB7t3-CR2<br>(EMDB:82007)<br>(PDB:43NF) |
| **Data collection and processing** |  |
| Microscope | FEI Titan Krios |
| Voltage (kV) | 300 |
| Detector | Gatan K3 Summit |
| Magnification | 130,000 × |
| Pixel size (Å) | 0.66 |
| Electron exposure (e<sup>-</sup>/Å<sup>2</sup>) | 50 |
| Defocus range (μm) | -1.2 to -2.0 |
| Automation software | EPU |
| Energy filter slit width (eV) | 20 |
| Movies used (no.) | 8,041 |
| Initial particle images (no.) | 1,887,172 |
| Final particle images (no.) | 384,769 |
| Symmetry imposed | C3 |
| Map resolution (Å) | 2.97 |
| FSC threshold | 0.143 |
| **Refinement** |  |
| Initial model used | AlphaFold |
| Model resolution (Å) | 3.0 |
| FSC threshold | 0.143 |
| Map sharpening *B* factor (Å<sup>2</sup>) | -153.1 |
| **Model composition** |  |
| Non-hydrogen atoms | 6201 |
| Protein residues | 819 |
| Ligands | 0 |
| **R.m.s.deviation**s |  |
| Bond lengths (Å) | 0.002 |
| Bond angles (°) | 0.471 |
| **Validation** |  |
| Molprobity score | 1.26 |
| Clash score | 5.01 |
| Rotamers outliers (%) | 0.00 |
| **Ramachandran plot** |  |
| Favored (%) | 99.63 |
| Allowed (%) | 0.37 |
| Outliers (%) | 0.00 |

